# The origin and evolution of archaeal Borg extrachromosomal elements

**DOI:** 10.64898/2026.05.22.727314

**Authors:** Ling-Dong Shi, Petar I. Penev, Bethany C. Kolody, Leylen Miloslavich, Shufei Lei, Rohan Sachdeva, Anna N. Rasmussen, Bradley B. Tolar, Christopher A. Francis, Alexander J. Probst, Xabier Vázquez-Campos, Timothy E. Payne, Zhou Jiang, Junxia Li, Cheng Wang, Zhili He, Jinren Ni, Laura A. Hug, Jillian F. Banfield

## Abstract

Borgs are giant linear extrachromosomal elements (ECEs) of methane-oxidizing *Methanoperedens* archaea whose evolutionary origin and ecosystem distribution remain unknown. Here we detected 240 Borgs in diverse saline, soil, and freshwater ecosystems. 27 encode methyl-coenzyme M reductases central to methane metabolism, eight have full rRNA operons related to those of *Methanoperedens*, and some contain up to four ribosomal proteins. We also identified 323 mini-Borgs whose proteome content and gene phylogenies classify as a distinct ECE type. Based on 105 complete and near-complete genomes, Borgs and mini-Borgs share core genes in conserved order, indicating common ancestry. Phylogenetic and diversity analyses suggest that Borgs evolved via gene acquisition into a backbone inherited from mini-Borgs. This evolutionary trajectory mirrors those proposed for giant viruses of both eukaryotes and bacteria.

## Main Text

Virtually all organisms harbor extrachromosomal genetic elements (ECEs) that impact host metabolism, cause cell death, and influence host evolution (*1–5*). The nature of host-ECE interactions is determined, in part, by ECE genome size (*6–8*). ECEs with large genomes have been reported (*9–12*), but their evolutionary trajectories remain controversial. It is argued that nucleocytoplasmic large DNA viruses (NCLDVs; *Nucleocytoviricota*) of Eukaryotes evolved from much smaller viral ancestors by capturing genetic material from their eukaryotic hosts, co-infecting bacteria, and other sympatric mobile elements (*13–15*). This may also be true for huge phages of Bacteria that are predicted to have evolved multiple times from smaller ancestral elements (*11*, *16–18*). Given the limited number of characterized large archaeal ECE genomes, it remains unknown whether this pattern applies to the evolution of large ECEs in Archaea. The recent discovery of giant Borgs provides an opportunity to address the question.

Borgs are linear ECEs associated with methane (CH_4_)-oxidizing *Methanoperedens* archaea (*12*). Their genomes currently range from 624 kbp to 1.095 Mbp in length and include prominent features such as long terminal inverted repeats (> 1 kbp), short genome-wide tandem repeats, and unidirectional gene coding on each replichore. Paralleling NCLDVs (*19*), Borgs integrate host-like pathways (e.g., translation apparatus and energy metabolism) that in NCLDVs are known to reprogram host cellular networks and drive eukaryotic adaptation to changing environments (*20–22*). Closely related to the Borgs is an evolutionarily distinct group of ECEs with identical genome architecture and some genes shared with Borgs; however, they have much smaller genomes (52-145 kbp) and thus are designated mini-Borgs (*23*). Both Borgs and mini-Borgs encode capsid-like proteins and show viral-like blooms in some samples (*19*, *22*, *23*), yet their classification as viruses has not been established in part due to the absence of canonical viral genes (e.g., terminase). Borgs and mini-Borgs have previously been reported from only five locations, and most genomes were reconstructed from a single wetland soil (*22*). Undersampling limits our understanding of Borg and mini-Borg environmental distribution, genomic diversity, gene repertoires, and evolutionary trajectories.

Here, we searched newly generated and publicly available metagenomes and found that Borgs and mini-Borgs occur in many diverse ecosystems. We invested significant manual effort in the curation of metagenomic assembled scaffolds to dramatically increase the number of complete genomes. This enabled detailed analyses of Borg and mini-Borg genome sizes, genome architectures, metabolic repertoires, and conserved, likely essential genes. Based on the expanded genome set, we propose an explanation for the origin and evolution of Borgs.

## Results

### A mini-Borg integrated into *Methanoperedens*

Prior analyses strongly indicated that *Methanoperedens* species host Borgs and mini-Borgs (*12*, *23*), but definitive evidence has been elusive. When searching for Borg and mini-Borg genes, we detected a 15 kbp element integrated in a circular *Methanoperedens* genome recovered from rice field soil. The element has 21 bp terminal inverted repeats, which are flanked by 14 bp direct repeats that define the integration site. The integrated region has ∼10% lower GC content compared to the *Methanoperedens* genome (35.1% versus 45.4%), as is typical for Borgs and mini-Borgs. The region bears short tandem repeats and encodes all genes on one DNA strand, features of the mini-Borgs with < 60 kbp genomes (Fig. S1) (*23*). Read mapping demonstrates that the integrated element has much lower abundance than the flanking host chromosome, and inconsistencies with read alignments to the consensus sequence confirm that the element is integrated into a subset of genomes of the *Methanoperedens* population. Of the 26 proteins encoded in the integrated element, two have homologs in *Methanoperedens* and five are most closely related to proteins of mini-Borgs (Figs. S2-S6). Based on these features, we conclude that the integrated element is a mini-Borg. This provides conclusive evidence that *Methanoperedens* is a host for mini-Borgs.

### Widespread distribution of Borgs and mini-Borgs

Given many indications that *Methanoperedens* are hosts for Borgs and mini-Borgs, we searched for these ECEs primarily in public sequence datasets that contain *Methanoperedens*, enabling us to broadly define their abundance, diversity, and ecosystem distribution (Fig. S7). *Methanoperedens* was identified in 5,011 metagenomes using Sandpiper (*24*), 2,372 of which had been assembled in the Logan project (Table S1) (*25*). Logan project assemblies and sequencing reads from the samples without assemblies were downloaded and queried for Borg and mini-Borg marker genes using HMMER (*26*) and GraftM (*27*), respectively. Metagenomes with significant hits were assembled locally for genome reconstruction and analyses. We also generated new metagenomes from samples collected from a wetland, a creek, and a rice paddy soil and searched the assemblies for sequences from Borgs and mini-Borgs (Table S2). In total, 240 Borg elements and 323 mini-Borg elements were identified, greatly expanding their known diversity (Fig. 1a). Borgs and mini-Borgs were detected at many sites across North America, Europe, and Asia (Fig. 1b and Table S2). Their habitats include groundwater, a drinking water system, municipal landfill, lake and river sediments, permafrost, grassland, and woodland soils. Notably, they also inhabit saline ecosystems, such as mangrove and marine sediments (Fig. 1a). Some sites had a very high diversity of Borgs and mini-Borgs. For example, 28 Borg species and 30 mini-Borg species were detected in a single groundwater sample (Fig. 1b).

**Figure 1.**
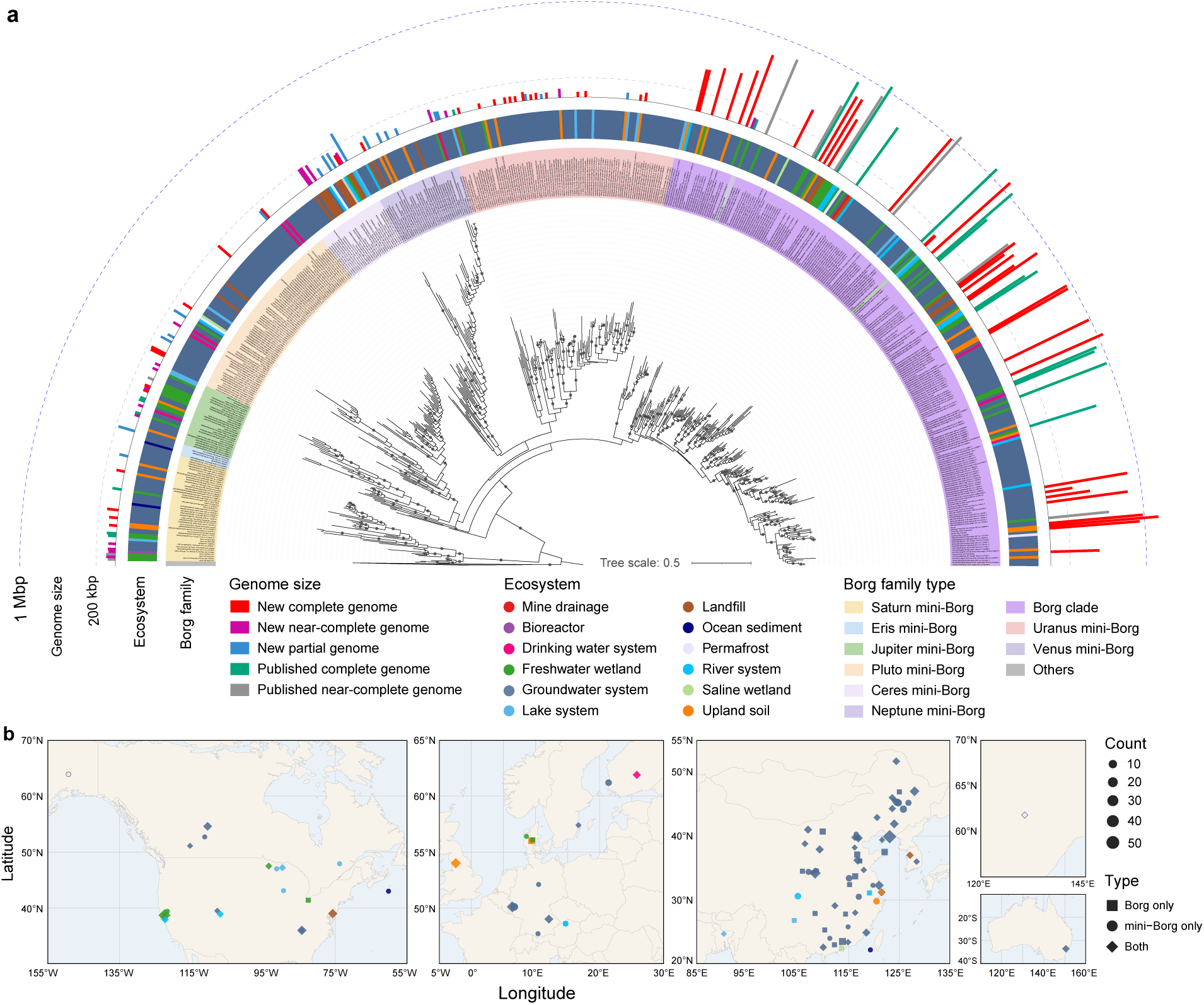
Distribution of diverse Borg family members across Earth’s ecosystems. **(a)** Phylogeny of Borgs and mini-Borgs based on a single-copy hypothetical protein (∼250 aa) using the best-fit substitution model “Q.yeast+R9”. The tree was rooted using non-Borg sequences as the outgroup. The inner to outer rings indicate Borg family genome type, environmental sampling type, and genome size. Branches are labeled when SH-aLRT values ≥ 80% and UFBoot values ≥ 95%. **(b)** Geographical map of samples with detectable Borg family members. Symbol colors indicate the ecosystem types of the samples, as described in (**a**). The sizes represent counts of Borg family members in the samples. The shapes denote the detectable presence of Borgs, mini-Borgs, or both.

The abundances of Borgs and mini-Borgs varied across samples, but their ratios compared to the sum of the abundances of all coexisting *Methanoperedens* were often very high, up to ∼300:1 (Fig. S8). They were even detected in ∼32% of samples in which *Methanoperedens* species were not detected. In some samples from wetland and upland soils, Borg and mini-Borg abundances exceeded the combined abundances of all bacteria and archaea (Fig. S8). The very high abundance ratios strongly support the possibility that Borg and mini-Borg DNA is present outside host cells and protected from degradation by encapsulation (*22*, *23*).

### Borgs and mini-Borgs as distinct ECE types

By extensive manual curation, we reconstructed 29 new, complete Borg genomes and 46 new, complete and near-complete mini-Borg genomes, tripling the available Borg genomes and increasing the number of mini-Borg genomes five-fold (Table S3). The new genomes substantially broaden the genome sizes for Borgs, which now range from 285 kbp (Mist Borg) to 1.146 Mbp (Mustard Borg), and for mini-Borgs, from 49 kbp (a Jupiter mini-Borg) to > 326 kbp (a Ceres mini-Borg) (Fig. 1a). The largest mini-Borg genome size now exceeds that of the smallest Borg.

The overlap in size ranges of mini-Borgs and small Borgs raised the possibility that the two ECE types form a single continuum. We identified 45 protein subfamilies that are encoded in > 90% of the 47 complete and near-complete Borg genomes as single copy, regardless of Borg genome sizes (Tables S4 and S5). Notably, 33 of these (73%) are present in Mist, the smallest Borg, yet only 16 of these (36%) are present in the largest mini-Borg (Fig. 2a). The count of these genes in mini-Borgs increases linearly with increasing genome size, yet there is a clear discontinuity in the trend for mini-Borgs and Borgs (Fig. 2a). Further supporting this indication that the two ECE types do not fall on a continuum, mini-Borgs are clearly separate from Borgs in the whole proteome profile (Fig. 2b).

**Figure 2.**
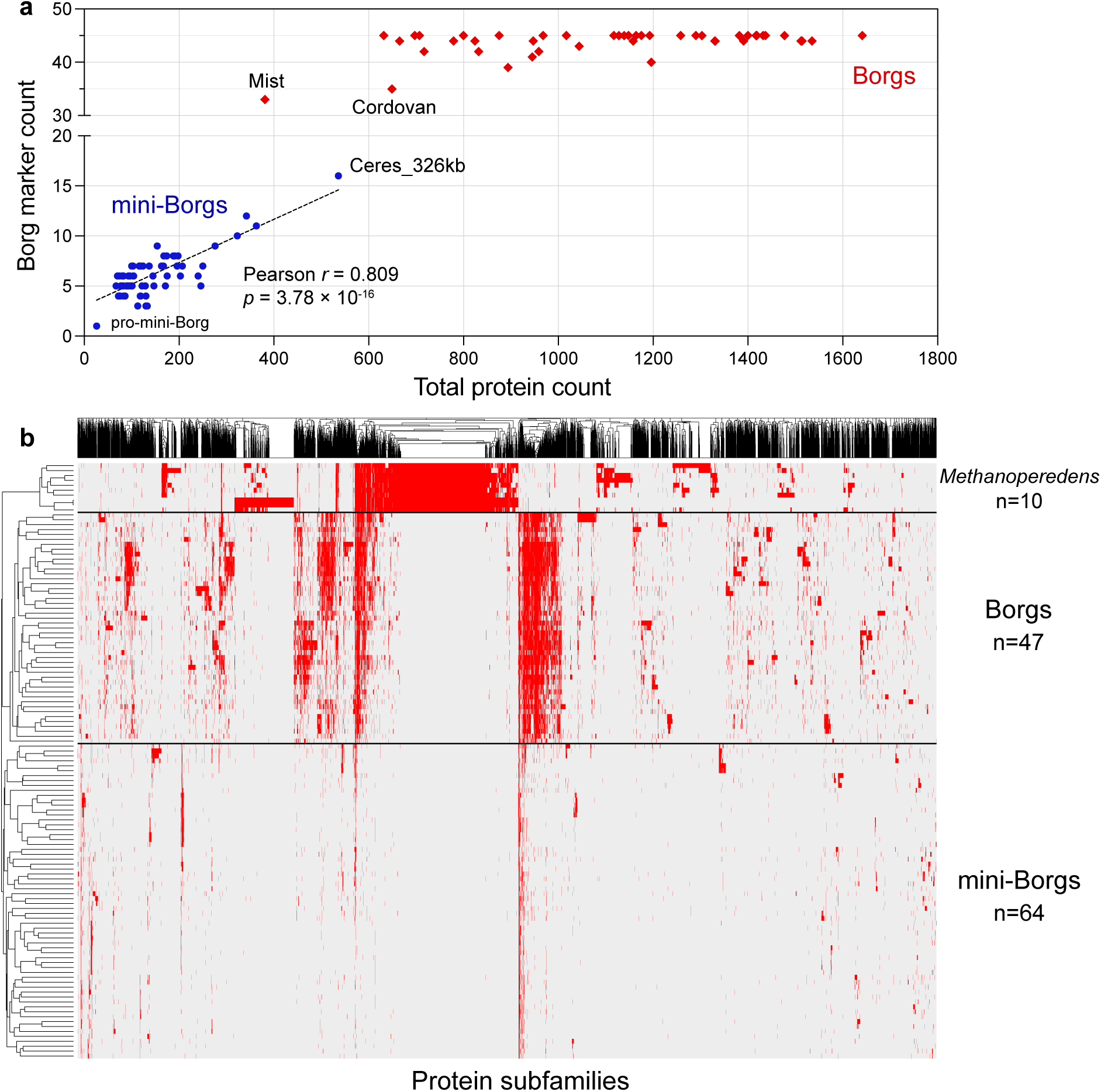
Proteome content of Borgs and mini-Borgs. (**a**) Count of 45 single-copy genes in Borg and mini-Borg genomes. The relationship of marker counts versus total protein counts in mini-Borgs was estimated by Pearson correlation (two-tailed test). Note the segmentation in the y-axis. (**b**) Distribution of protein subfamilies across Borgs, mini-Borgs, and *Methanoperedens*. Genomes were clustered based on the presence (red)/absence (gray) profiles using Jaccard distance (complete linkage).

### Expanded Borg gene repertoires central to CH_4_ metabolism

Borgs encode inventories of key metabolic genes central to CH_4_ metabolism (*12*). Here we identified methyl-coenzyme M reductase complex (MCR) in three complete Borg genomes and on 24 Borg scaffolds from groundwater, drinking water, lake sediment, upland and wetland soil (Fig. 3a). Borg-encoded McrA sequences nest within the *Methanoperedens* clade, with indications of multiple lateral gene transfer events from *Methanoperedens* hosts to Borgs. Intriguingly, most Borgs encode *mcrBDGA* in operons, as do *Methanoperedens*, but nine Borg gene clusters lack *mcrD*. Flanking the Borg *mcr* operons, and in some cases between *mcr* genes, are short tandem repeats (TRs) (Fig. S9). TRs are a notable feature of Borg genomes but do not occur in proximity to, or within, *Methanoperedens mcr* operons. This provides confidence that the *mcr* operons are not misassembled into Borg genomes.

**Figure 3.**
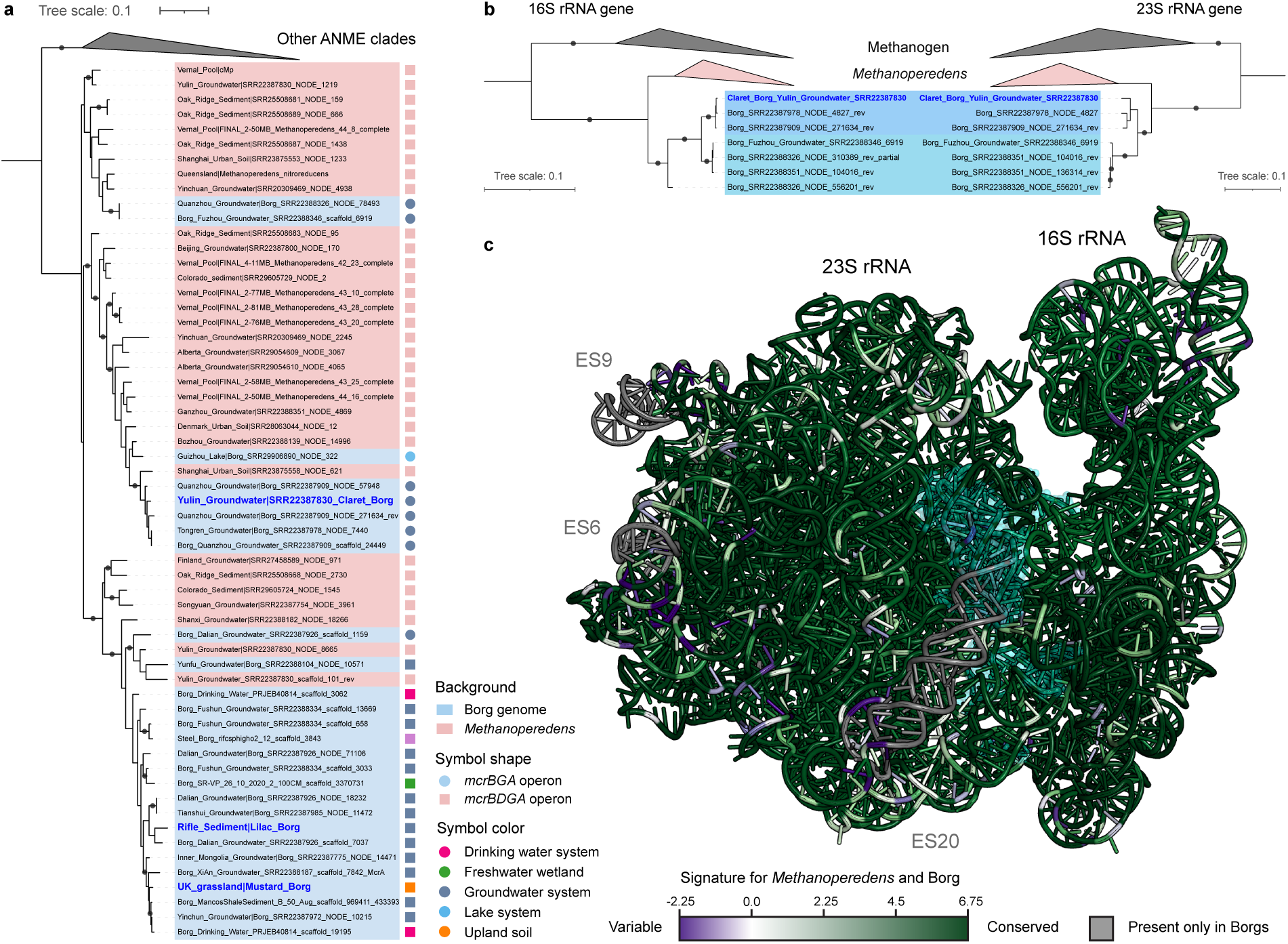
Examples of Borg organismal genes. **(a)** Phylogeny of McrA subunit using the best-fit substitution model “LG+F+R4”. Sequences in the blue background are from Borg genomes and those in the red background are from *Methanoperedens*. Borg genomes in blue and bold font are curated and complete. Genomes labeled by circles encode *mcrBDGA* operons and those labeled by squares encode *mcrBGA* operons. Symbol colors indicate ecosystems wherein Borgs existed. (**b**) Phylogeny of 16S and 23S rRNA genes using the best-fit substitution model “GTR+F+I+R2”. Sequences in the blue background are from a curated, complete Claret Borg and additional, incomplete Borg genomes. Note that “Borg_SRR22388326_NODE_310389_rev_partial” has too low coverage to recover the intact 23S rRNA gene, thus it is not included in the 23S rRNA gene tree. “Borg_SRR22388351_NODE_136314_rev” contains the partial 16S rRNA gene and therefore is not included in the 16S rRNA gene tree. Branches are labeled when SH-aLRT values ≥ 80% and UFBoot values ≥ 95%. **(c)** Predicted structures of Borg rRNAs colored by conservation between Borg and *Methanoperedens* sequences. Conserved nucleotides between Borg and *Methanoperedens* are in dark green, conserved but deferring nucleotides are in purple, and nucleotides with low overall conservation are in white. Nucleotides absent from *Methanoperedens* but present in the Borg rRNAs are in gray. The residues interacting between the two rRNAs are shown with cyan transparent spheres.

MCR activates CH_4_, converting heterodisulfide to methyl-coenzyme M and coenzyme B. Heterodisulfide is replenished by the heterodisulfide reductase complex (HDR) (*28*). We identified a fused, membrane-bound *hdrDE* gene in Coffee Borg that is homologous to sequences in some *Methanoperedens* genomes. The predicted structure aligns well with the separate HdrD and HdrE subunits from *Methanosarcina* (Fig. S10). Additionally, cytoplasmic *hdrABC* operons were also found in Coffee Borg and on four other Borg scaffolds. The Borg HdrA sequences were distributed in two phylogenetic clades, members of which have *Methanoperedens* homologs (Fig. S11a). Most Borg *hdrABC* showed sequence deviation compared to host genes (71.9 ± 1.58% nucleotide identity, Fig. S11b), but the operon in Coffee Borg (∼3,800 bp) shared up to 96.7% nucleotide identity with various *Methanoperedens* operons, suggesting recent gene transfer (Fig. S11c).

### Ribosomal RNAs and proteins encoded in Borgs

Surprisingly, eight Borg genomes comprising one subclade that includes the curated, complete Claret Borg, encode full rRNA operons (16S rRNA, tRNA, 23S rRNA, and 5S rRNA genes). The 16S and 23S rRNA gene phylogenies show congruent topologies, with Borg sequences nested within those from *Methanoperedens* (Fig. 3b). The average pairwise identity of 16S rRNA genes is 97.6% within Borgs and 93.9% between Borgs and *Methanoperedens*, indicating an approximately family-level separation of the two groups (Fig. S12) (*29*).

We modeled the structures of Borg rRNAs and found that all regions critical for translation are present, indicating that they are functional (Figs. 3c and S13-S14). Compared to the *Methanoperedens* rRNA structure, the modeled Borg 23S rRNA structure differs in regions that primarily associate with the ribosome E-site, as well as some regions associated with the central protuberance and the 5S rRNA binding (Figs. 3c and S13). There are three expansion segments (ES) in the Borg 23S rRNAs, positioned in the canonical locations of ES6, ES9, and ES20 (*30–32*). ES20 is the largest (60 nucleotides) and is modeled to form a long helical structure (Fig. 3c).

We modeled the ribosomal positions of the Borg 16S and 23S rRNAs based on a superimposition to the complete ribosomal structure of *Pyrococcus furiosus* (PDB: 4V6U). The two Borg rRNAs show no major conflicts (Fig. 3c). On the small subunit, many of the differences between Borg and *Methanoperedens* rRNAs are in the head region (Fig. 3c), suggesting changes in the decoding and proofreading process of translation. The interface between the large and small subunits is well conserved between *Methanoperedens* and Borg rRNAs (Fig. 3c). This indicates that likely the two subunits can interact with no changes in *Methanoperedens* ribosomes.

Borgs collectively encode eight different ribosomal proteins, most of which are components of the small ribosomal subunit (Fig. S15). Coffee and Burgundy Borg genomes each encode four ribosomal proteins, including consecutive rpS13-rpS4-rpS11 that share up to 87% nucleotide identity with syntenous sequences in coexisting *Methanoperedens* genomes (Table S5). The rpS4 and rpS11 proteins bind the small subunit far from the decoding center of the ribosome. The rpS13 protein has a C-terminus that models as an alpha helix with an extension that is long enough to reach the ribosome decoding center.

### Phylogeny and evolution of Borgs and mini-Borgs

Mini-Borgs share only six near-universal single-copy genes, three of which are in the 45 conserved Borg gene set (Table S6). These three protein sequences were aligned and concatenated to generate a Borg and mini-Borg phylogeny in which all Borgs cluster together, although three mini-Borgs are placed within the Borg clade with strong branch support (Fig. 4a). The Borg clade topology is consistent with that in the tree constructed by concatenating the 45 Borg gene markers (Fig. S16), and in the larger phylogeny containing partial genomes (Fig. 1a). All other mini-Borgs cluster within six clades (e.g., Jupiter) that are distinct from Borgs, and each clade exhibits larger sequence diversity than that of Borgs (Fig. 4a).

**Figure 4.**
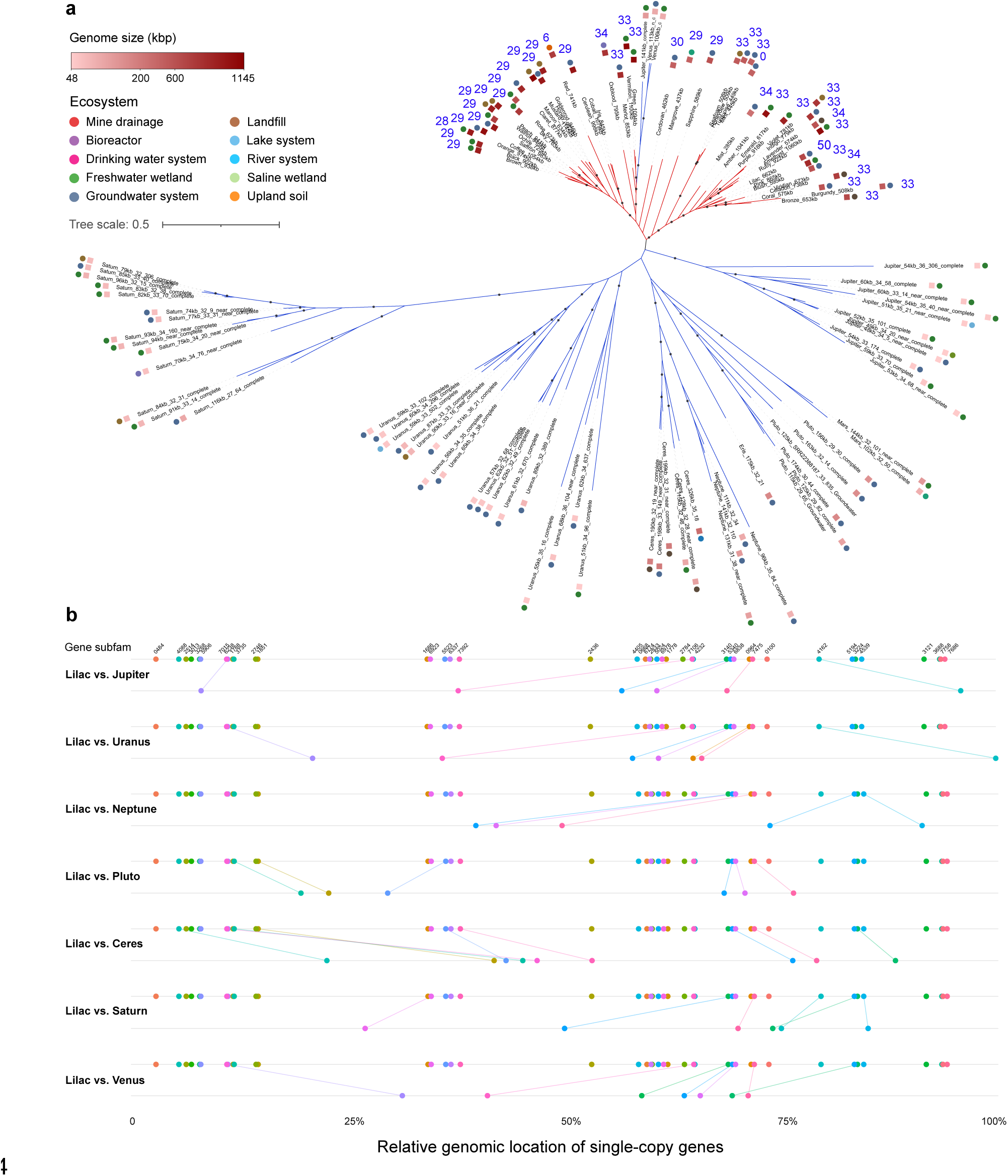
Phylogeny and synteny of essentially complete Borg and mini-Borg genomes. (**a**) The phylogenetic tree is built based on 3 concatenated, single-copy proteins (∼500 aa) present in Borgs (red branches) and mini-Borgs (blue branches) using the best-fit substitution model “LG+I+R5”. Branches are labeled when SH-aLRT values ≥ 80% and UFBoot values ≥ 95%. Rectangle colors next to clade names indicate genome sizes. Circles are colored by ecosystems from which genomes were recovered. Numbers in blue denote the count of the 58 mini-Borgs that share significantly conserved orders of single-copy genes with Borgs. (**b**) Syntenic localization of single-copy marker genes on Borg and mini-Borg genomes. Dots indicate the relative genomic locations of the marker genes normalized by total protein counts. Lines connect the same genes between pairs of genomes. Lilac Borg is compared with representative mini-Borgs from each clade.

Different mini-Borgs encode different subsets of the 45 single-copy genes (Fig. 2a), but the subsets and counts of these genes are largely conserved in each mini-Borg clade (Fig. S17). Notably, within each mini-Borg clade, the order of these genes is also conserved.

We wondered if genes present in both Borgs and mini-Borgs occur in the same order, consistent with inheritance of a genomic backbone from a common ancestor. As it is well established that Borgs share a conserved backbone (*19*, *22*), we developed a metric for synteny that was thresholded using Borg gene order conservation scores (Fig. S18). For each Borg, we counted the number of mini-Borgs that exceeded the threshold score indicative of gene order conservation. Lilac Borg has the largest number of mini-Borgs with significant conservation of order of shared single-copy genes (50 of the 58 mini-Borgs with (near-)complete genomes, and from all clades; Fig. 4). Generally, Borgs closely related to Lilac exhibit synteny with more mini-Borgs than distantly related Borgs. Interestingly, Mint Borg encodes most of these genes in reverse order compared to other Borgs and mini-Borgs (Figs. 4a and S18). This is due to a large genome rearrangement anchored at the terminus of replication that switched the small and large replichores and the associated unidirectional coding (Fig. S19). Overall, the analyses are indicative of a backbone in Borgs and mini-Borgs that was inherited from a common ancestor.

To attempt to resolve whether Borgs evolved into mini-Borgs or vice versa, we analyzed the phylogeny of the 19 single-copy genes that are encoded in ≥ 10 genomes of both groups. Borg and mini-Borg sequences intermix in nine of these trees (Figs. S20-S28), indicative of non-uniform evolutionary trajectories (e.g., horizontal gene transfer). Thus, these trees are not informative regarding origination. In six trees, Borgs and mini-Borgs form distinct, separate clades (Figs. S2 and S29-S33). Two of these six trees could be rooted using an outgroup (S32-S33), yet in both trees the Borgs and mini-Borgs are placed as independent clades with no indication of evolutionary direction. In the remaining four trees, one ECE type is nested within the other (Figs. 5a and S34-S36). These four genes are distributed unevenly in mini-Borgs but are universal in Borgs, indicating that Borgs acquired them before further diversifying. After rooting these four trees using microbial sequences as outgroups, Borg sequences nest within mini-Borg sequences (Figs. 5a and S34-S36), suggesting that Borgs inherited these four genes from mini-Borgs. Also suggesting gene inheritance from mini-Borgs, the B9 protein-primed DNA polymerase (PolB9), which occurs in a Neptune mini-Borg (but no other mini-Borgs), is basal to the Borg clade (Fig. S37). PolB9 is encoded in all Borg genomes very early in the large replichore and is likely essential for genome replication (*19*). In summary, gene phylogeny suggests that Borgs evolved from mini-Borgs.

**Figure 5.**
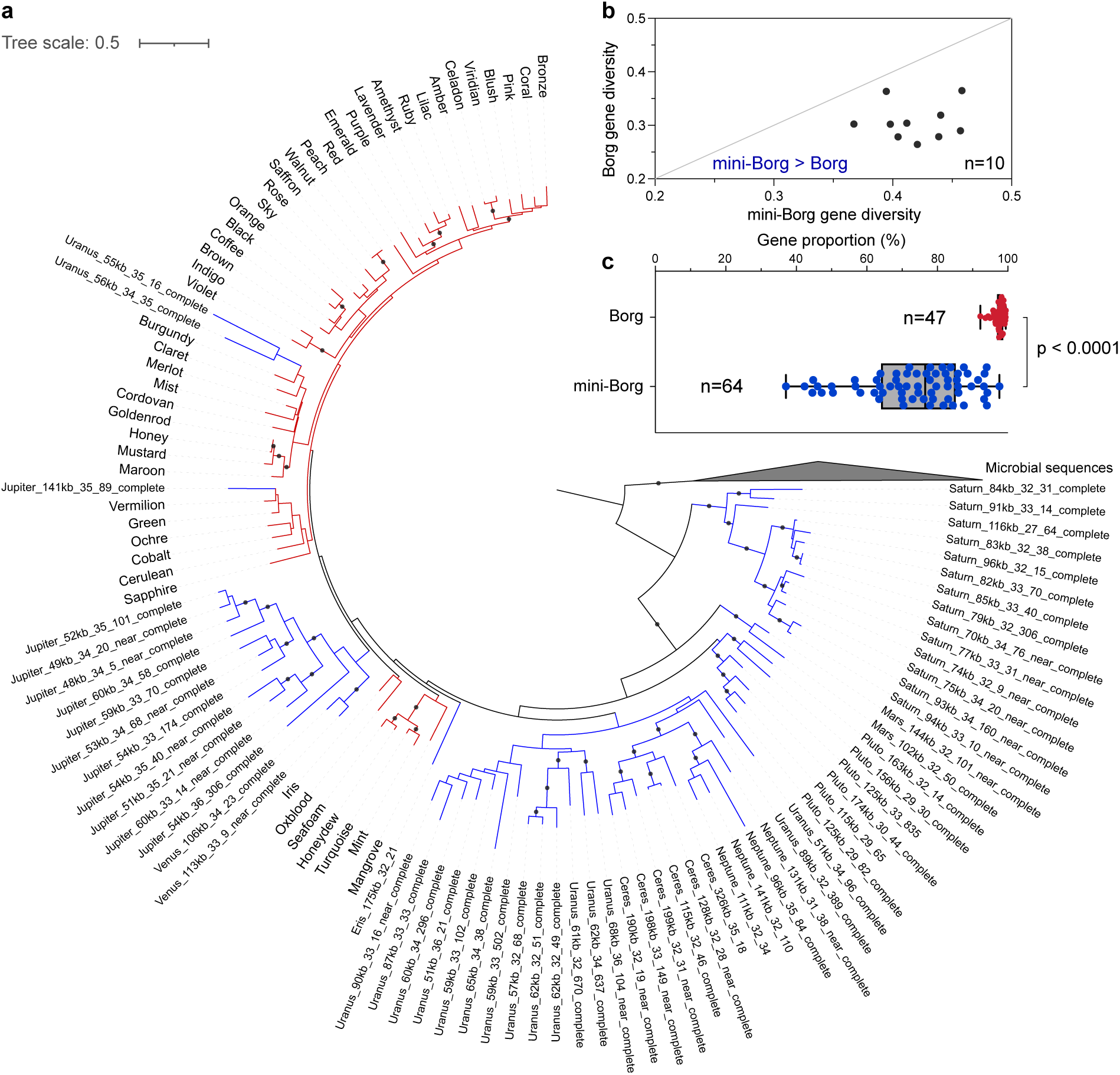
Genetic evidence inferring Borgs evolved from mini-Borgs. **(a)** Phylogeny of single-copy putative proteases (subfam7475) encoded in Borgs and mini-Borgs using the best-fit substitution model “LG+F+I+R5”. Branches in red and blue indicate Borg and mini-Borg genes. The tree was re-rooted using microbial sequences as the outgroup. Branches are labeled when SH-aLRT values ≥ 80% and UFBoot values ≥ 95%. Note that all Borgs and mini-Borgs encode this protein. (**b**) Nucleotide diversity of 10 single-copy genes shared by Borgs and mini-Borgs. Borg and mini-Borg sequences generally form distinct phylogenetic clusters that separate from one another. Detailed phylogenetic trees of individual genes are present in Figs. 5a, S2, and S29-S36. (**c**) Proportion of genes best matching the ingroup sequences. The statistical difference between the two groups was evaluated using Welch’s Two-Sample t-test.

If Borgs arose from mini-Borgs, we anticipate that they will display less genetic diversity than mini-Borgs. To test this, we calculated the nucleotide-level sequence diversity for the 10 of the 19 proteins that did not show evidence of extensive lateral gene transfer (Figs. 5a, S2 and S29-S36). The mini-Borgs display significantly higher genetic diversity than Borgs in all cases (Fig. 5b). Supporting this conclusion, 97.6 ± 1.5% of Borg proteins have the best-matching homologs in another Borg, whereas only 73.8 ± 14.7% of mini-Borg proteins were identified in more than one mini-Borg (Fig. 5c).

Borg and mini-Borg genome sizes vary substantially within clades, even between the most closely related genomes, suggestive of relatively recent gene gain and/or loss (Fig. 4a). For example, the Blush and Amber Borg genomes share 81% average nucleotide identity (ANI) over 59% of the Blush genome, yet the Blush genome is only 57% of the size of Amber. Similarly, two Ceres mini-Borgs share 69% ANI over 33% of the smaller genome, but the Ceres_115kb_32_46 genome size is only 58% of Ceres_199kb_32_31. We infer that genome size change is ongoing throughout both lineages.

## Discussion

We analyzed the public sequence read archive and newly generated metagenomic data, a total of > 5,000 metagenome datasets comprising ∼90 Tbp of sequence information, to document the environmental distribution and genomic diversity of Borgs and mini-Borgs (Table S1). This was an important undertaking given that, to date, these ECEs have only been reported in publications by one laboratory. We estimated that these ECEs occurred in metagenomes generated by at least 39 different research groups (Table S2). From a subset of these data we manually curated 58 ECE genomes to completion and 17 to very near completion. We used the expanded set of ultra-high-quality genomes to address the question of the relatedness of Borgs to mini-Borgs. Mini-Borgs were named due to features shared with Borgs, including their genome architectures, overlap in gene content, and *Methanoperedens* host-association, but their genome sizes were much smaller (*23*). Although one newly reported (yet incomplete) Ceres mini-Borg genome is larger than the smallest complete Mist Borg genome, gene phylogenies and whole proteome content distinguish mini-Borgs from Borgs unambiguously (Figs. 2 and 4). Thus, we conclude that Borgs and mini-Borgs are distinct. Borgs and mini-Borgs share some DNA modification patterns when they occur in the same sample (*33*), suggesting that they can coexist in the same *Methanoperedens* cell, enabling genetic exchange with each other as well as with their hosts.

Extending knowledge of their environmental distribution in wetland soils, groundwater, and freshwater sediment, we found Borgs and mini-Borgs in diverse ecosystems. In some cases, Borgs and mini-Borgs are the most abundant genomic components in their microbiomes (Figs. 1 and S8). Given that they can account for a significant fraction of microbiome DNA, they likely have the potential to regulate their host and overall microbiome function across many of Earth’s environments. The possible ecological significance of Borgs and mini-Borgs could derive from their impacts on host activity, either via the energetic cost of their maintenance and replication or the provision of beneficial genes, or both.

Most notable of the Borg genes are those with the potential to modulate host CH_4_ metabolism. As a methanotrophic archaeon, the host *Methanoperedens* conserves energy primarily from anaerobic CH_4_ oxidation (*34*). Central to this process is the MCR complex, an enzyme encoded in CH_4_-metabolizing archaea and predicted to originate from the last Archaeal common ancestor (*35*, *36*). Previously, the MCR operon was reported in only one complete Borg genome and on one Borg scaffold (*12*), but not in any other ECEs. Here, we identified MCR complexes in 25 additional Borg genomes from geographically distinct ecosystems associated with active CH_4_ cycling (Fig. 3a) (*37–39*). Notably, some Borgs encode *mcrBGA* gene operons that differ from the *mcrBDGA* in *Methanoperedens* but still share high sequence similarity with host genes. The missing gene *mcrD* is proposed to facilitate the assembly of the cofactor F_430_ into the MCR complex (*40*). However, this gene is dispensable in methanogens and is absent in ANME-1 methanotrophs (*40*, *41*). Borgs likely acquired the full operon from *Methanoperedens* (ANME-2d) and subsequently lost the D subunit. By duplicating gene content, Borgs may partially mitigate CH_4_ emissions via promotion of host CH_4_ oxidation. The availability of a likely functional but sequence distinct MCR complex, possibly with modified enzyme kinetics, may extend the conditions under which methane metabolism can occur. Analogous to phage therapy, we speculate that, long term, Borgs could be introduced into environments hospitable to *Methanoperedens* to reduce emissions from CH_4_ hotspot ecosystems.

It is argued that encoding ribosomes defines cellular organisms (*42*), as even the smallest intracellular parasites encode a ribosome, including rRNAs (*43*, *44*). ECEs rarely encode rRNAs. The only known non-chromosomal elements with rRNA operons are small bacterial plasmids (< 53 kbp) (*45–47*). Here, we show that one group of Borgs encodes full rRNA operons that were horizontally acquired from the host *Methanoperedens* (Fig. 3b). The structural analysis indicates that the Borg rRNAs are functional (Fig. 3c). The Borg 23S rRNAs contain three expansion segments (ES) that do not occur in the *Methanoperedens* sequences (Fig. 3c). ESs are considered a momentary snapshot of ribosomal evolution following the accretion model (*48–50*). The existence of variable ES regions suggests that the Borg 23S rRNAs are subject to evolutionary pressure to expand. This may contribute to the specialization of some *Methanoperedens* ribosomes, as observed in eukaryotes (*51*, *52*). Given the 93.9% pairwise identity of 16S rRNA genes between Borgs and *Methanoperedens*, and an estimated 16S rRNA divergence rate of 1% per 50 million years (*53*, *54*), the subclade of Borgs with rRNAs may have been co-occurring with *Methanoperedens* for > 300 million years.

As Borgs do not encode the full set of ribosomal proteins, they cannot be considered to be organisms. The ribosomal proteins identified in Borgs are S1, S2, S4, S7, S11, S12, S13, and L11 (Fig. S15). This suggests that the Borgs with rRNAs hijack host ribosomal proteins for building hybrid *Methanoperedens*-Borg ribosomes. This is reminiscent of mitochondria and chloroplasts in eukaryotes, which utilize their own rRNAs and nuclear ribosomal proteins to synthesize ribosomes for translation of their encoded genes (*55*, *56*). Some Borgs encode sequential rpS13, rpS4, and rpS11 that may be co-transcribed. The extension on the C-terminus of rpS13 is predicted to reach the decoding center, where mRNA translation, codon-anticodon interactions and mRNA translocation occur during protein biosynthesis (*57*). We suspect that Borg proteins may assemble into hybrid ribosomes to alter mRNA binding and select for Borg transcripts preferentially over host transcripts.

Variable subsets of the 45 fairly syntenous single-copy genes that are distributed throughout the Borg genomes are also encoded in mini-Borg clades, where they occur with a highly conserved order (Figs. 4 and S17). The shared backbone strongly indicates common ancestry. Borgs may have streamlined their genomes and evolved into mini-Borgs, so mini-Borgs represent newer lineages. According to molecular clock theory (*58*), Borgs then should have accumulated greater genetic variation than mini-Borgs, similar to the higher diversity observed in human African mitochondrial DNA versus non-African DNA (*59*). However, all genes in mini-Borgs show significantly higher nucleotide diversity than those in Borgs (Fig. 5). Phylogenomic analysis of shared core genes revealed that mini-Borgs are paraphyletic, yet Borgs cluster in a single monophyletic clade (Fig. 4a). These findings support an alternative, parsimonious model that Borgs evolved from mini-Borgs by vertically inheriting the genome backbone and laterally acquiring genes from hosts and other sources. The evolution of mini-Borgs into Borgs is further supported by individual gene phylogenies, which nest Borg sequences within those of mini-Borgs in cases where root positions can be confidently inferred (Figs. 5a and S34-S37). Expanded genome size enabled Borgs to accommodate a vast repertoire of auxiliary metabolic genes, some of which were acquired recently from *Methanoperedens* hosts (e.g., MCR and HDR). Due to smaller genome sizes, mini-Borgs (even the largest Ceres mini-Borg) are mostly constrained to core, non-host genes.

From where the mini-Borg originated remains unknown. The discovery of a 15 kbp pro-mini-Borg in the *Methanoperedens* genome may provide some insight, given that it shares identical genome architecture and conserved genes with mini-Borgs (Fig. S1). This is reminiscent of Polintons (or Mavericks), which are integrated in eukaryotes, have typically 15-20 kbp genomes, contain terminal inverted repeats, are flanked by direct repeats, and encode several conserved proteins shared with viruses (*60–62*). They are hypothesized to have evolved from a bacterial tectivirus-like ancestor and given rise to various large eukaryotic DNA viruses (*14*, *63*, *64*). Analogous to the evolutionary relationship between Polintons and eukaryotic viruses, the element integrated in the *Methanoperedens* genome may represent the progenitor of mini-Borgs, yet more examples are needed to explore this hypothesis.

In conclusion, Borgs carry a versatile gene repertoire, sometimes including components of ribosomes and gene operons central to host methane metabolism. Thus, they may modulate *Methanoperedens* metabolic capacity, transcriptional activity, and ecological fitness. Our findings indicate that Borgs evolved from mini-Borgs via the acquisition of genes throughout their genomes. The evolutionary trajectory of archaeal Borgs well aligns with that observed in the giant viruses of eukaryotes and bacteria (*11*, *13–18*), suggesting that the process of genome expansion leads to ECE complexity across all three domains of life.

## Materials and Methods

### Sample collection and processing

Soil samples were collected from a wetland site at a depth of 50 - 60 cm in Lake County and from a rice field at a depth of 20 - 30 cm in Butte County, California, USA. Water samples were collected from Strawberry Creek at the UCB campus by filtering water through 0.1 µm polyethersulfone filters to retain microbial biomass. Samples were frozen immediately using liquid nitrogen for DNA extraction. DNA was extracted using Qiagen DNeasy PowerMax Soil Kit and sequenced on the Illumina platform. The rice paddy soil sample was also sequenced using the PacBio HiFi long-read technology. Short reads were processed by trimming or removing low-quality sequences using BBDuk (https://jgi.doe.gov/data-and-tools/software-tools/bbtools/) and assembled using metaSPAdes (v4.2.0) (*65*). Long reads were assembled using metaMDBG (v0.3) (*66*).

### Search for Borgs and mini-Borgs

To explore Borgs and mini-Borgs in public datasets, we leveraged Sandpiper (*24*), which has screened 707,470 metagenomes and profiled community structure, to search for the presence of their host *Methanoperedens* by the broader taxonomy “f Methanoperedenaceae”. This yielded 5,011 samples containing detectable *Methanoperedens* as of May 21, 2025 (Fig. S1). We mapped sample accession numbers to those in Logan (*25*) and obtained 2,372 metagenomic assemblies. For 2,639 samples that were absent in Logan, we downloaded raw reads from NCBI and investigated the presence of Borgs and mini-Borgs using GraftM with a custom gene package (*27*). The package was created based on a ∼250 aa protein (subfam7475, see below) that is universal across all available Borgs and mini-Borgs (*23*). We then processed and assembled reads from samples containing Borgs and mini-Borgs, as described above, generating the assembly set that includes new samples and public datasets (Fig. S1).

Genes were predicted from assemblies using Prodigal (v2.6.3) with the DNA code 11 (*67*), and compared with previously identified Borg marker genes using BLASTp (v2.16.0+) (*22*, *68*). Potential Borg and mini-Borg scaffolds were recovered if they encode marker genes and have GC contents of 30-35% (*12*, *23*).

### Genome curation

Illumina reads were mapped to potential Borg and mini-Borg scaffolds with 5% mismatches to check and fix assembly errors, including N gaps, indels, inconsistent single nucleotide polymorphisms (SNPs), and collapsed repeats, following our previous paper (*69*). As Borgs and mini-Borgs are enriched in tandem short repeats, the collapse of repeats is a common issue caused by assemblers. This error was identified by finding reads with discrepant regions. We first identified and labeled repeats in reads, and then searched for discrepant read regions and determined if that sequence is part of a repeat. The placement of the discrepant region outside of the repeat region likely indicated that the repeat region was collapsed. We then opened a gap in the assembly and added the missed sequence in repeats, consistent with the largest number of repeats on any read. In cases where the reads indicated a variable number of repeat units in the population, the largest repeat unit count supported by the data was generally chosen.

For a clean scaffold without errors, the Illumina reads placed at scaffold ends and their unplaced paired reads were used to extend the sequences. The extended sequences sometimes merged with other scaffolds based on overlap. This process was repeated until no further extension was possible, either because the genome was complete and inverted repeats had been identified, or because there were too few reads to extend. Then reads were mapped to refined scaffolds with 0% error to check if the entire scaffold was covered by paired reads evenly. For regions with particularly low coverage under these circumstances, reads were mapped less stringently to find better consensus sequences. In cases of low overall coverage, a 1% mismatch was allowed for final mapping to check the quality and accuracy of curated genomes. Final genomes were subjected to pairwise comparison using the online blastn program.

### Gene annotation

Gene sequences were compared against references in the KEGG (release 114.0) (*70*) and UniProt databases (release 2025_01) by USEARCH (v10) (*71*), and those in the NCBI nr database (release 257.0) by BLASTp (v2.16.0+) (*68*). Protein domains were identified by HMMER3 (v3.3) (*26*) using the Pfam 37.0 database with pre-defined Noise Cutoff (NC) bit score thresholds (*72*).

Structures of protein monomers were predicted by AlphaFold2 via LocalColabFold (v1.5.5) with default parameters (*73*, *74*). Structures of multimers were modeled on the AlphaFold3 web server (*75*). Monomeric folds labeled “relaxed_rank_001” and having median pLDDT scores ≥ 70 were compared against structures in the PDB database using Foldseek (version 427df8a6b5d0ef78bee0f98cd3e6faaca18f172d) (*76*). Only best matches with *e*-values ≤ 0.001, bit scores ≥ 50, and query and target coverages ≥ 50% were retained. Comparison of protein structures was performed and visualized in ChimeraX (v1.7.1) (*77*). Annotations based on sequences and structures were integrated and cross-validated for each protein.

### Environmental distribution of Borgs and mini-Borgs

To explore the presence and diversity of Borgs and mini-Borgs, a single-copy and universal protein (subfam7475) was extracted from all complete and near-complete genomes, based on which a custom Hidden Markov model (HMM) was created using HMMER3 (v3.3) (*26*). We performed an HMM-based search on metagenome samples and collected proteins that contain the same domain with high significance. Protein sequences were aligned with MAFFT using the parameters “--maxiterate 1000 --localpair” (v7.453) (*78*). The alignment was trimmed by trimAl “-gappyout” (v1.4.rev15) (*79*) and used to construct the phylogenetic tree by IQ-TREE (v2.3.6) with automatically selected best-fit models using ModelFinder “-m MFP” (*80*, *81*). The whole-tree robustness was evaluated using ultrafast bootstrap approximation (UFBoot) with 1,000 replicates (“-B 1000”). Branch support was assessed using the Shimodaira-Hasegawa approximate likelihood ratio test (SH-aLRT) with 1,000 replicates (“-alrt 1000”). The tree was formatted using the iTOL web server (*82*). Clades of Borgs and mini-Borgs were determined based on the tree topology.

### Abundances of Borgs, mini-Borgs, and prokaryotes

Abundances of scaffolds that contain the single-copy subfam7475 gene (∼250 aa) were used to indicate abundances of Borg and mini-Borg species. Abundances of scaffolds that contain the single-copy ribosomal protein S3 (rpS3, ∼250 aa) were used to indicate abundances of prokaryotic species. The subfam7475 gene was identified as described above. The rpS3 gene was identified using HMMER3 with the PF00189 and preliminarily classified by blasting against the NCBI nr database. Precise taxonomy was assigned to sequences by analyzing their phylogeny with references, following the protocol in the preceding section.

Scaffolds containing targeted genes were dereplicated at approximately species-level 97% identity. Metagenomic reads were cross-mapped to scaffolds using CoverM v0.6.1 (*83*).

The “contig” mode was used with a minimum read identity of 97% and a minimum aligned percent of 90%. If a scaffold had < 50% of bases covered by reads, it was regarded as absent in samples.

### Structural analysis of Borg rRNAs

The secondary structures of the Borg rRNAs were modeled on the R2DT web server (*84*) and adjusted with the online Exornata (*85*). The tertiary structures were modeled using the AlphaFold3 web server (*75*). Base-pairing of the new ES helices was determined based on the 3D modeled structures and extracted by ModeRNA (v1.7.5) (*86*). Conservation and divergence between the *Methanoperedens* and Borg rRNAs were calculated with TwinCons (v0.6.2) (*87*) using the nucleotide blastn matrix. The large and small subunits of the Borg ribosome were oriented by superimposing to the archaeal *Pyrococcus furiosus* ribosomal structure (PDB: 4V6U) (*88*) and visualized using PyMOL.

### Identification and localization of Borg single-copy marker genes

Proteins of 47 complete and near-complete Borg genomes and 64 complete and near-complete mini-Borg genomes were clustered into subfamilies using the method described previously (*89*). A protein subfamily encoded as a single copy and in ≥ 90% of Borgs is designated as a Borg single-copy marker gene. In total, 45 Borg single-copy marker genes were identified, some of which were encoded in mini-Borgs (Table S4).

Given the huge difference in genome length, we calculated the relative localization of Borg single-copy marker genes in Borg and mini-Borg genomes by normalizing with total gene counts. Ordinal association of marker genes between genome pairs was measured using the Kendall rank correlation test.

### Phylogeny and nucleotide diversity of Borg single-copy marker genes

Single-copy marker genes shared by ≥ 10 Borgs and ≥ 10 mini-Borgs were blasted against the NCBI nr database (release 257.0) and the GTDB database (r220) (*90*). We aligned query sequences with the recruited references, trimmed the alignments, and constructed gene phylogenies using the same protocol as above.

Nucleotide sequences of marker genes that are phylogenetically separate between Borgs and mini-Borgs were aligned and trimmed. Nucleotide diversity was calculated for Borg genes and mini-Borg genes using the R package pegas (v1.4) (*91*).

## Acknowledgements

The authors thank Dr. Luis E. Valentin-Alvarado for contributions to data collection and analysis, Dr. Tomas Hessler for assistance with bioreactor enrichment, and Dr. Shuai Wang and Dr. Ming Yan for constructive comments. We also thank the many investigators worldwide who generated datasets submitted to the SRA.

## Funding

The Emerson Collective via the Innovative Genomics Institute (JFB)

The Chan Zuckerberg Initiative award CZIF2022-007203 via the Innovative Genomics Institute (JFB)

U.S. Department of Energy, Office of Science, Office of Basic Energy Sciences, Chemical Sciences, Geosciences, and Biosciences Division, through its Geoscience program at LBNL, DE-AC02-05CH11231 (JFB)

The Gates Foundation grant INV-037174 (JFB)

Lawrence Livermore National Laboratory Microbes Persist SFA, DOE award SCW1632 (JFB)

The Lawrence Berkeley National Laboratory Watershed SFA (JFB)

Lyda Hill Philanthropies, Acton Family Giving, the Valhalla Foundation, Hastings/Quillin Fund, the CH Foundation, Laura and Gary Lauder and Family, the Sea Grape Foundation, the Emerson Collective, Mike Schroepfer and Erin Hoffman Family Fund, the Anne Wojcicki Foundation (RS)

The Southern Marine Science and Engineering Guangdong Laboratory (Zhuhai) grants SML2024SP002, SML2024SP022, SML2023SP205 (CW, ZH)

The National Natural Science Foundation of China grant 42430707 (ZH) A Canada Research Chair (LAH)

U.S. Department of Energy, Office of Science, Office of Biological and Environmental Research, Climate and Environmental Sciences Division, through its support of the former SLAC Floodplain Hydro-Biogeochemistry SFA Award Number DE-AC02-76SF00515, and Subsurface Biogeochemistry Program (SBR) Project Award Number DE-SC0016544 (CAF)

DOE Joint Genome institute, a DOE Office of Science User Facility supported by the Office of Science of the U.S. Department of Energy under Contract No. DE-AC02-05CH11231, supported some sequencing of samples through a Community Science Program grant #1927 (CAF) and Facilities Integrating Collaborations for User Science grant #504298 (BBT, CAF)

German Research Foundation (DFG), project PR 1603/4-1 (AJP)

## Author contributions

Conceptualization: LDS, JFB

Data generation: LDS, BCK, ANR, BBT, CAF, AJP, XVC, TEP, ZJ, JL, CW, ZH, JN, LAH

Genome curation: LDS, JFB

Gene annotation and analysis: LDS

Environmental distribution and abundance quantification: LDS

Structural analysis: LDS, PIP

Genomic backbone identification: LDS, JFB Phylogenetics and gene diversity calculation: LDS Bioinformatic support: LM, SL, RS

Writing – original draft: LDS, JFB

Writing – review & editing: All the authors.

## Data availability

The newly generated metagenomic sequencing data are available under NCBI BioProject PRJNA1463535. The genomes described in this study can be accessed at https://ggkbase.berkeley.edu/Borg_family_members.

## Supplementary Materials

Figs. S1 to S37

Tables S1 to S6

