## Supplementary Figures for "The origin and evolution of archaeal Borg extrachromosomal elements"

**The PDF file includes:**

Figures S1 to S37

**Other Supplementary Materials for this manuscript include the following:**

Table S1 to S6

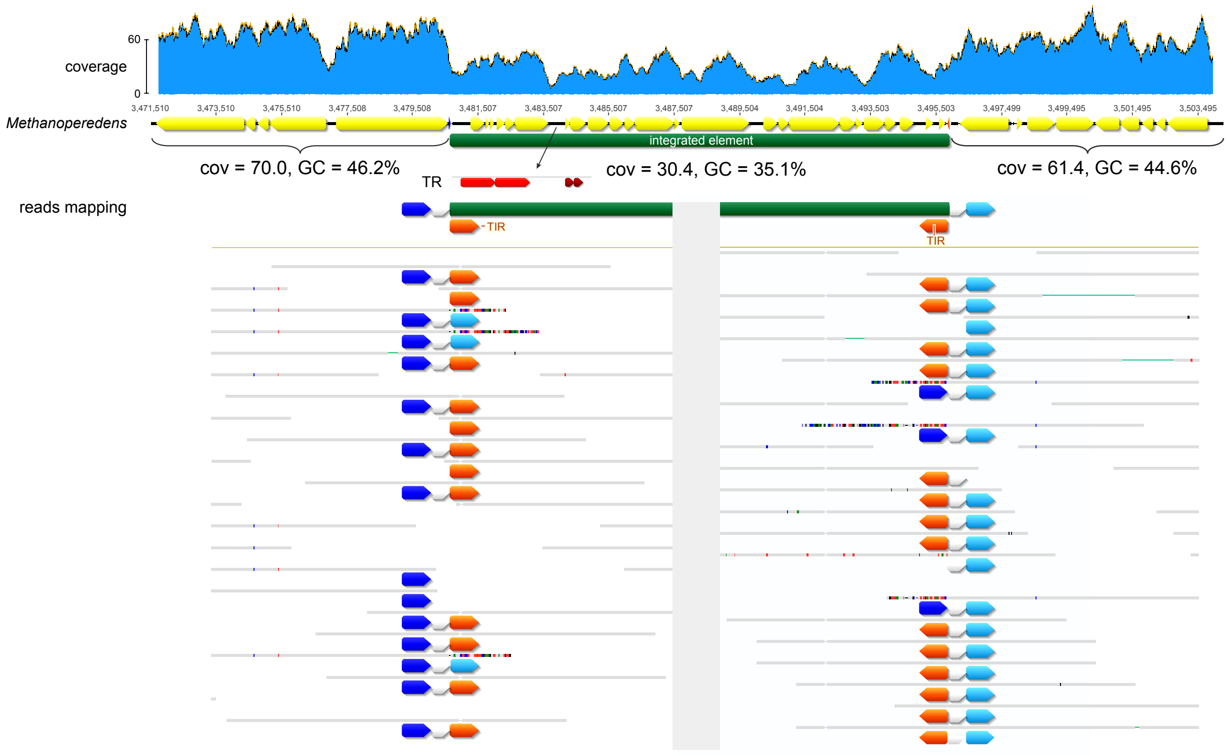


**Figure S1. An integrated element in the *Methanoperedens* chromosome (160_C).** Gray bars indicate reads that match the genome. Colored parts inside reads are segments that do not match the reference. Arrows with the same colors indicate identical sequences. Abbreviations: TR, tandem repeats; TIR, terminal inverted repeats.

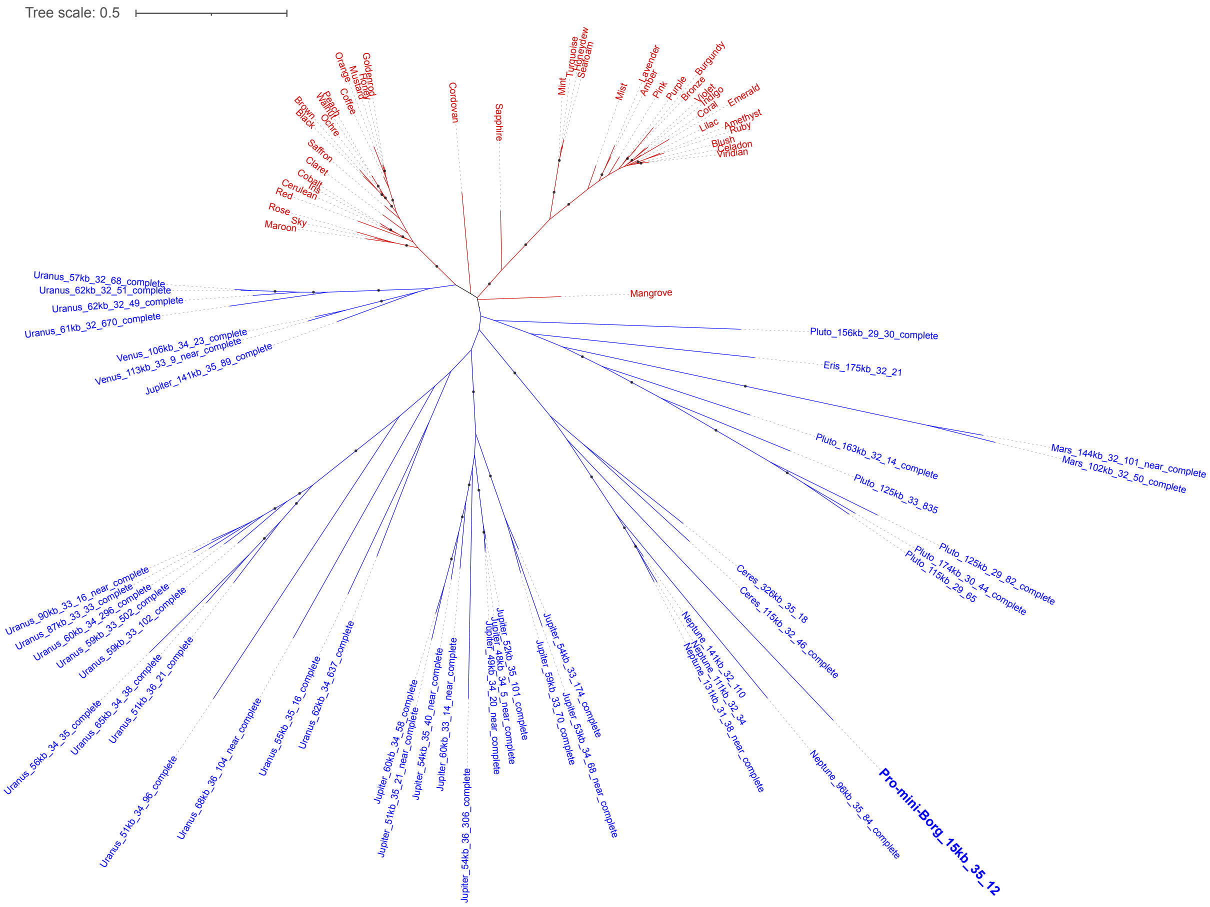


**Figure S2. Phylogeny of single-copy hypothetical proteins (subfam6838) encoded in the Borgs and mini-Borgs.** Sequences (branches) in red and blue indicate Borg and mini-Borg genes. The bold blue gene is encoded in the integrated mini-Borg. No homologs are found in other elements. The tree was constructed using the best-fit substitution model “Q.yeast+F+I+R4”. Branches are labeled when SH-aLRT values ≥ 80% and UFBoot values ≥ 95%.

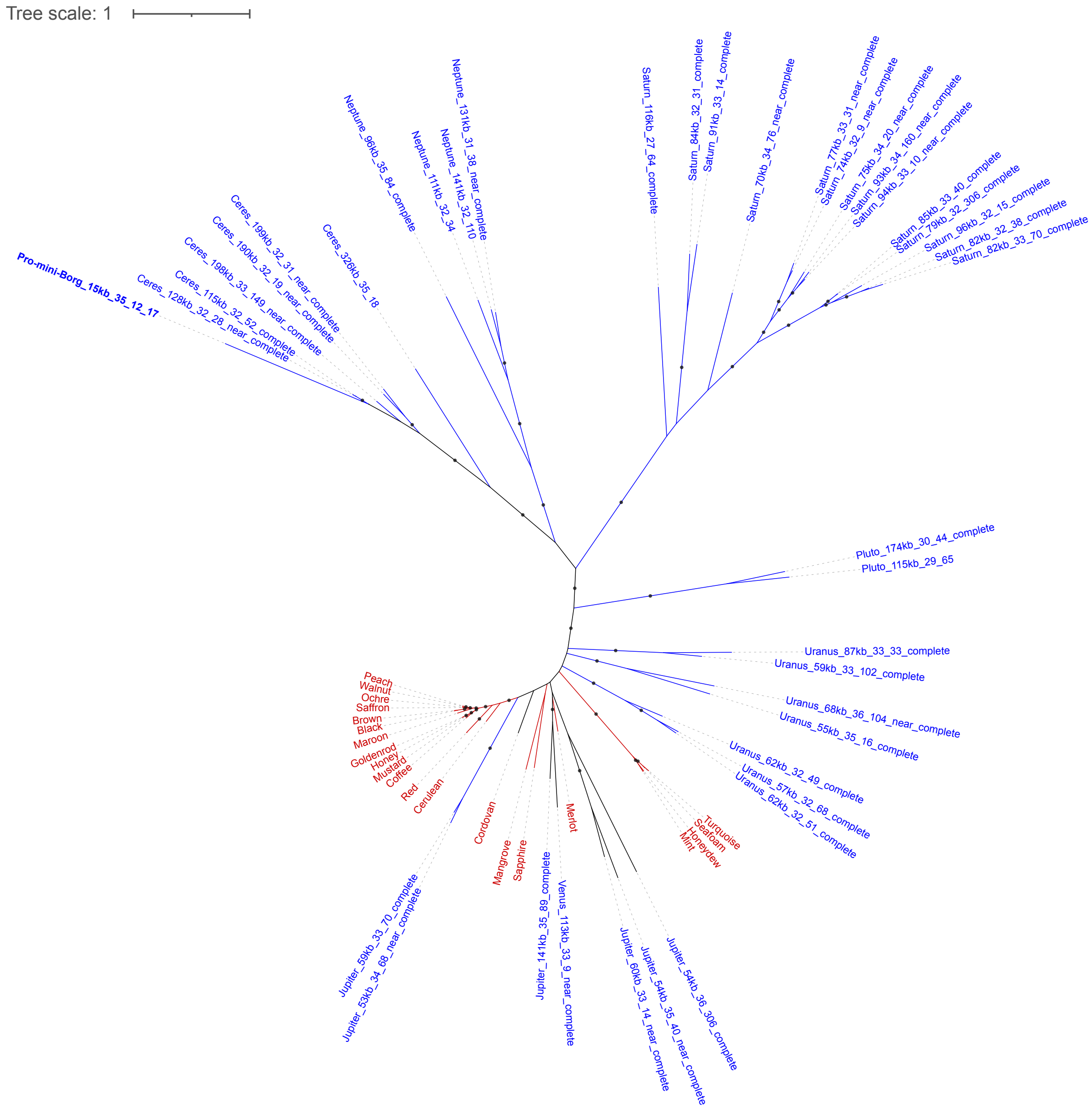


**Figure S3. Phylogeny of the integrated mini-Borg protein (ID_17).** Sequences (branches) in red and blue indicate Borg and mini-Borg genes. The bold blue gene is encoded in the integrated mini-Borg. No homologs are found in other elements. The tree was constructed using the best-fit substitution model "Q.insect+F+R5". Branches are labeled when SH-aLRT values ≥ 80% and UFBoot values ≥ 95%.

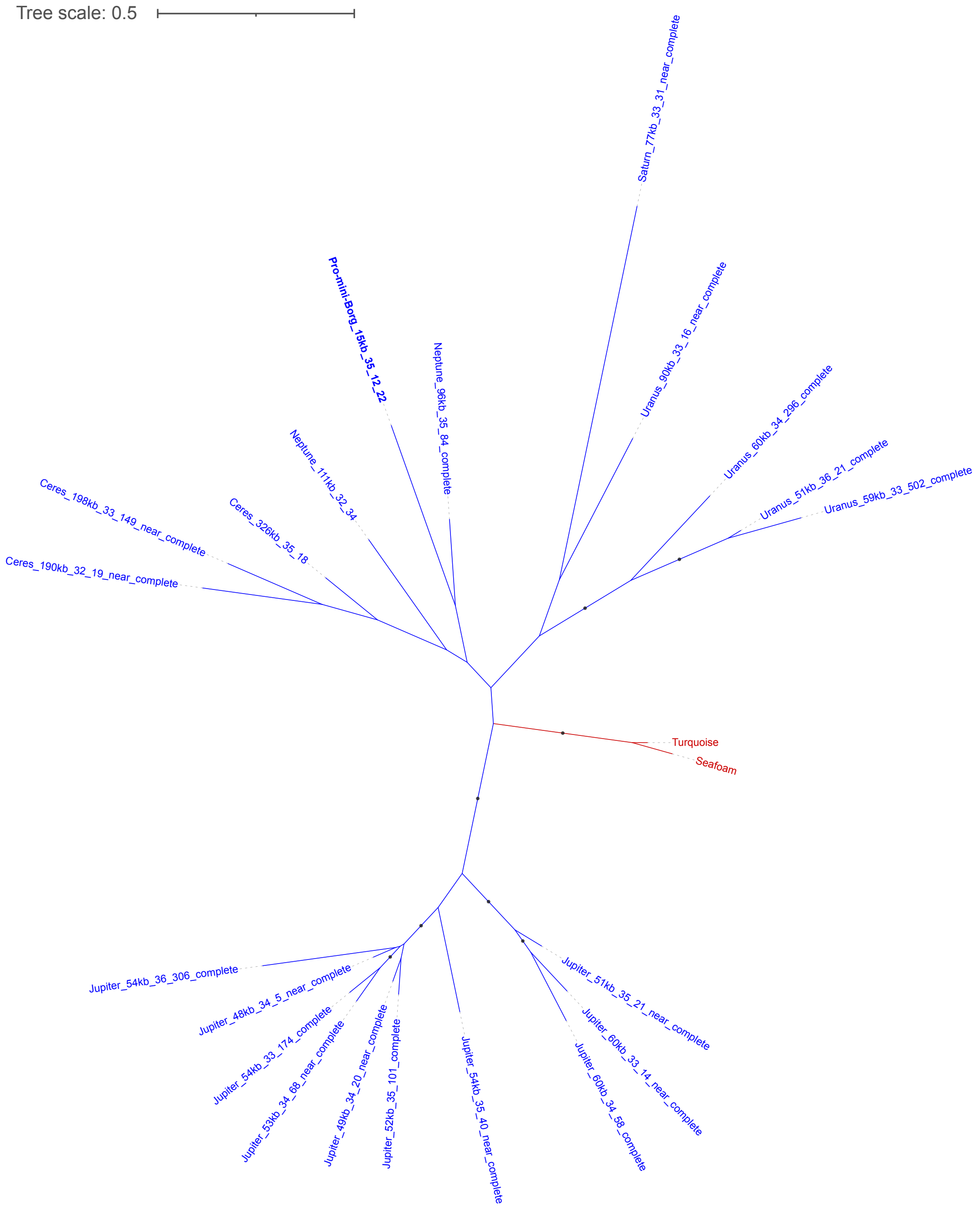


**Figure S4. Phylogeny of the integrated mini-Borg protein (ID_22).** Sequences (branches) in red and blue indicate Borg and mini-Borg genes. The bold blue gene is encoded in the integrated mini-Borg. No homologs are found in other elements. The tree was constructed using the best-fit substitution model “VT+I+R2”. Branches are labeled when SH-aLRT values ≥ 80% and UFBoot values ≥ 95%.

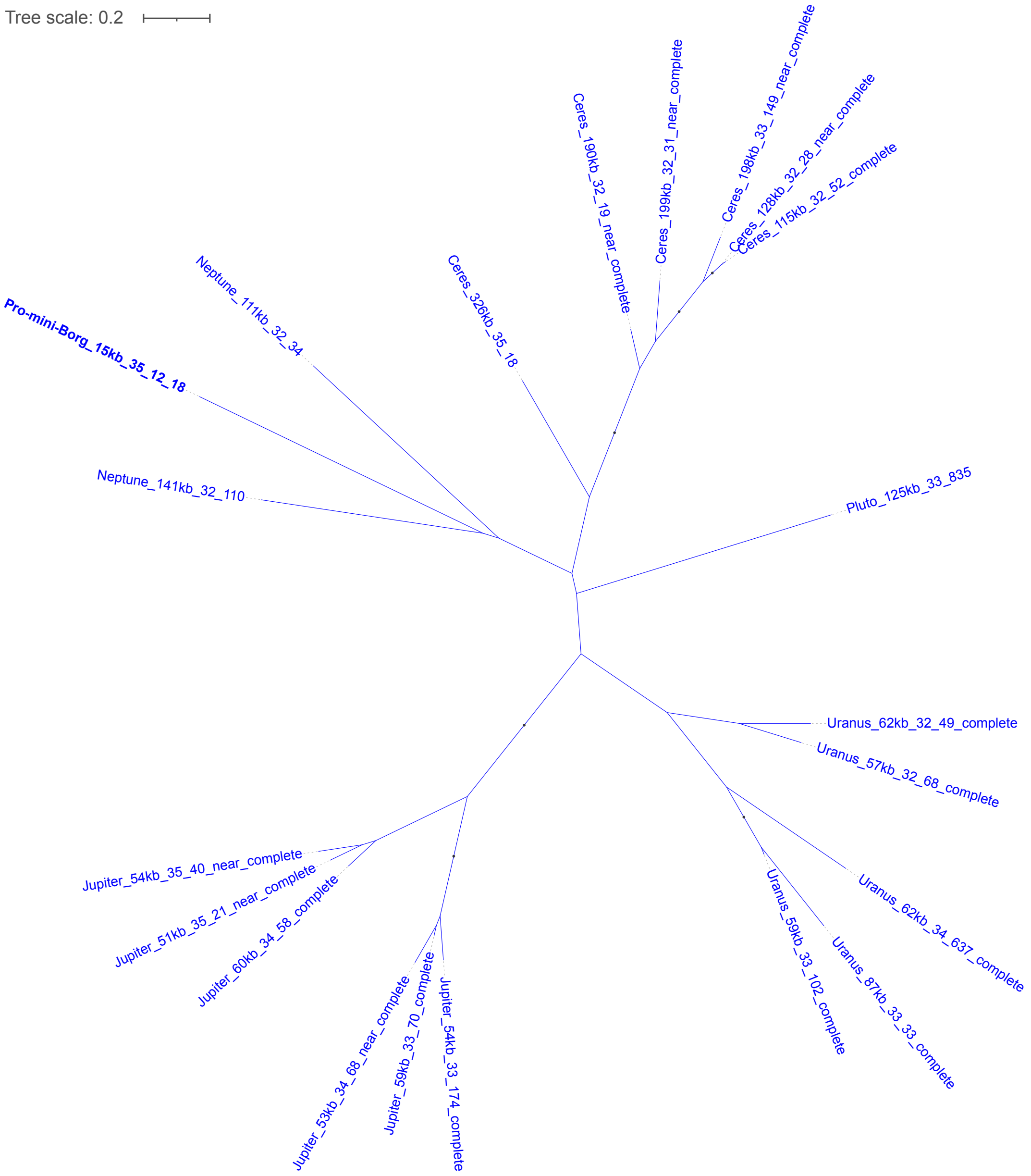


**Figure S5. Phylogeny of the integrated mini-Borg protein (ID_18).** Homologs are only found in mini-Borgs. The bold blue gene is encoded in the integrated mini-Borg. The tree was constructed using the best-fit substitution model "WAG+F+I+G4". Branches are labeled when SH-aLRT values ≥ 80% and UFBoot values ≥ 95%.

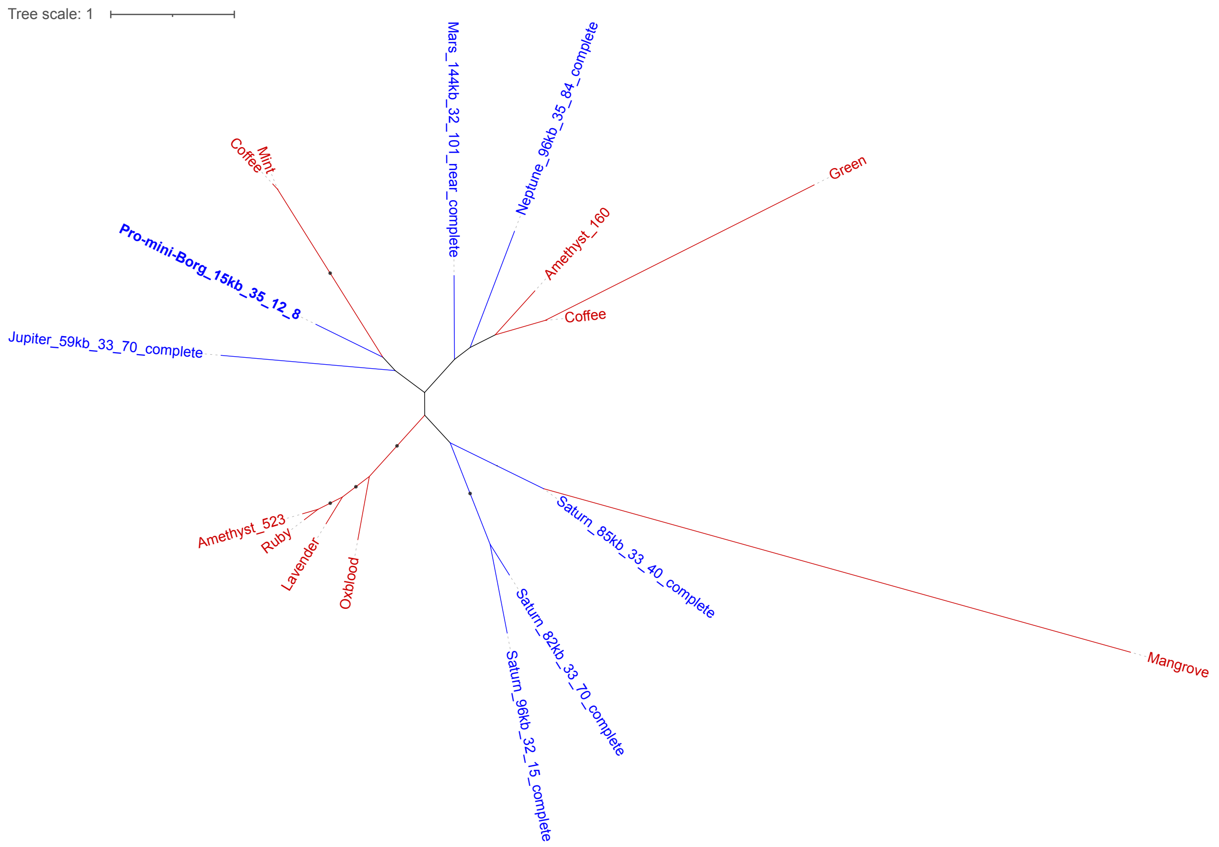


**Figure S6. Phylogeny of the integrated mini-Borg protein (ID_8).** Sequences (branches) in red and blue indicate Borg and mini-Borg genes. The bold blue gene is encoded in the integrated mini-Borg. No homologs are found in other elements. The tree was constructed using the best-fit substitution model “VT+F+G4”. Branches are labeled when SH-aLRT values ≥ 80% and UFBoot values ≥ 95%.

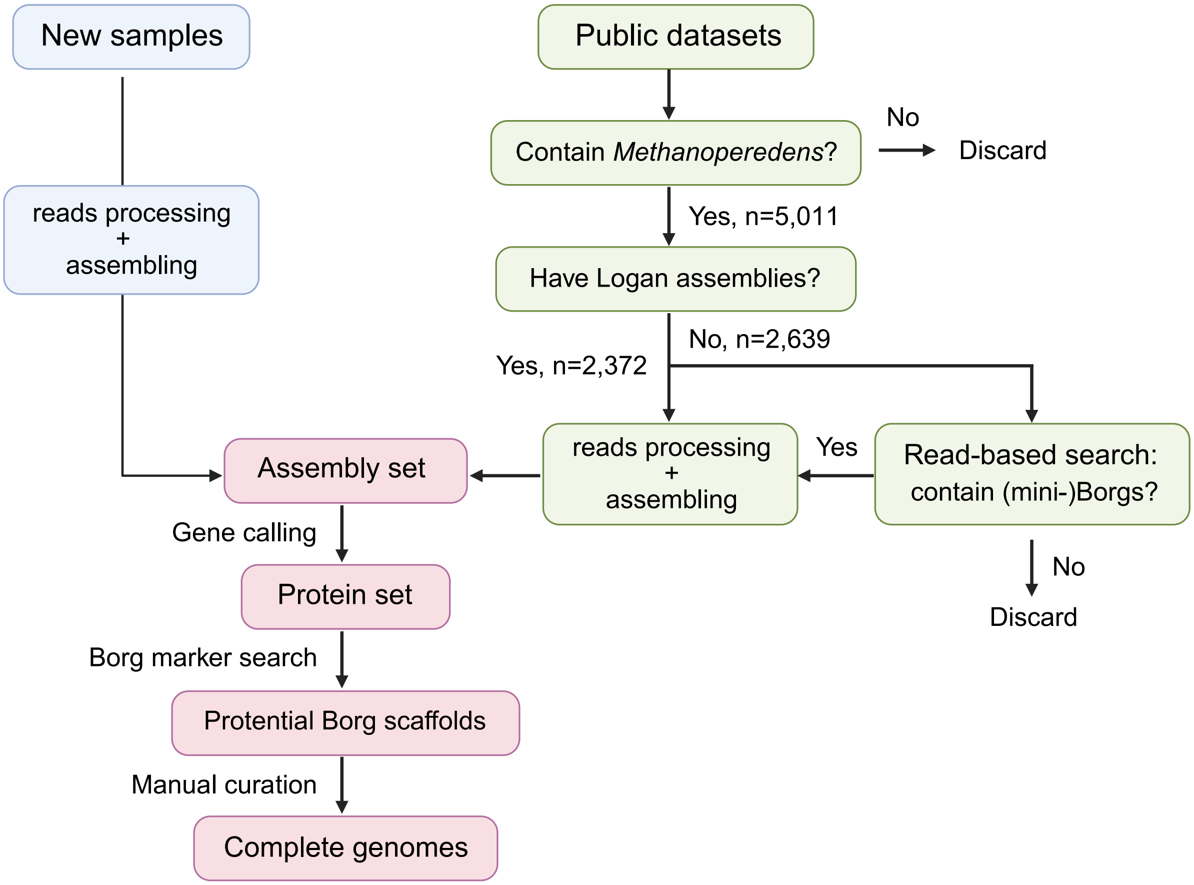


**Figure S7. Workflow of Borg search in metagenome samples.** Details can be found in Materials and Methods.

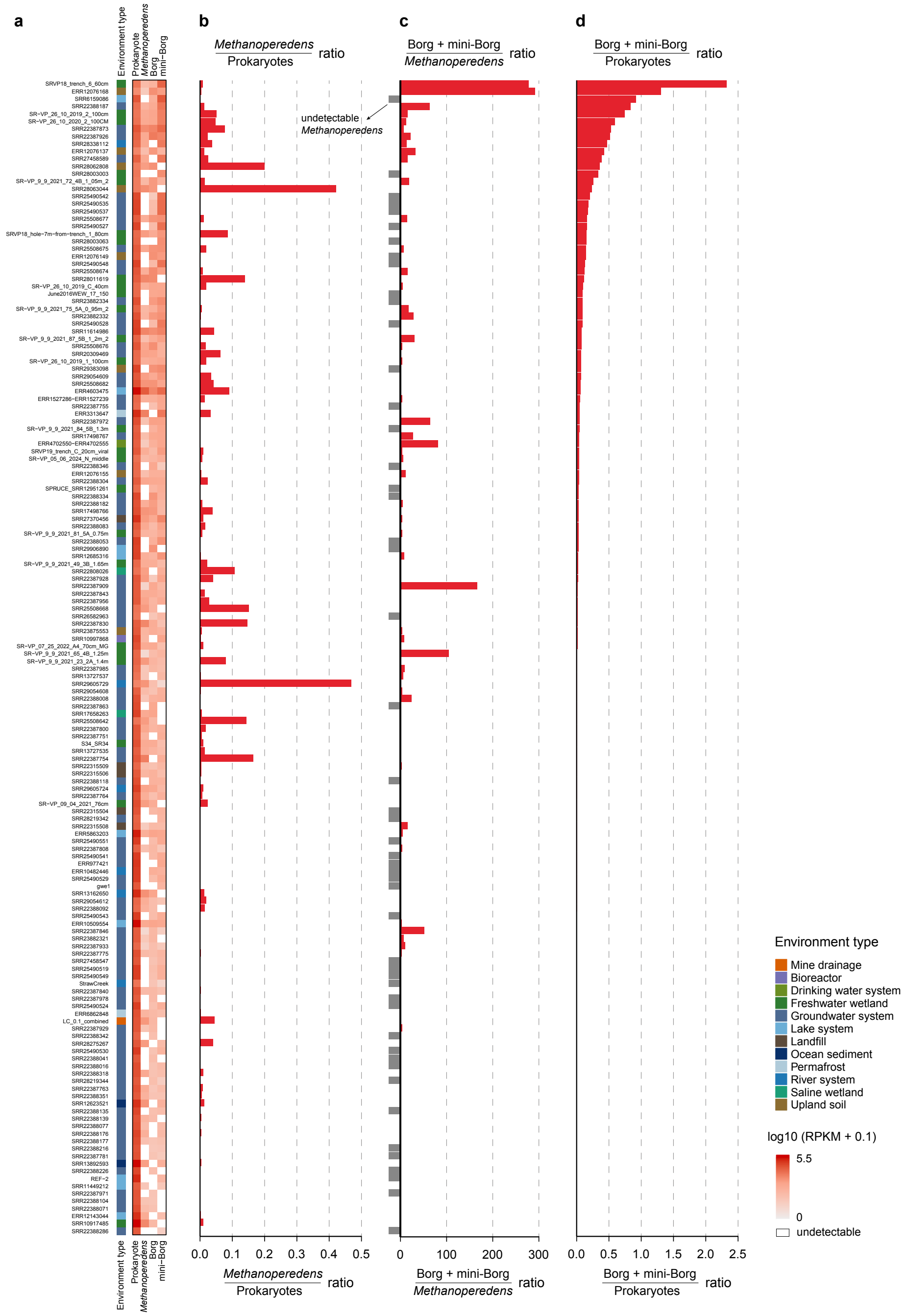


**Figure S8. Distribution of Borgs, mini-Borgs, and *Methanoperedens* in the prokaryotic community across samples. (a)** Sum abundances of Borgs, mini-Borgs, *Methanoperedens*, and the entire prokaryotic community. Abundances are converted logarithmically for visualization purposes. Blank indicates undetectable presence. **(b)** Abundance ratios of *Methanoperedens* in the prokaryotic community. **(c)** Abundance ratios of Borgs and mini-Borgs to *Methanoperedens*. Minus values (gray bars) indicate the undetectable existence of *Methanoperedens* in samples. **(d)** Abundance ratios of Borgs and mini-Borgs to all prokaryotes.

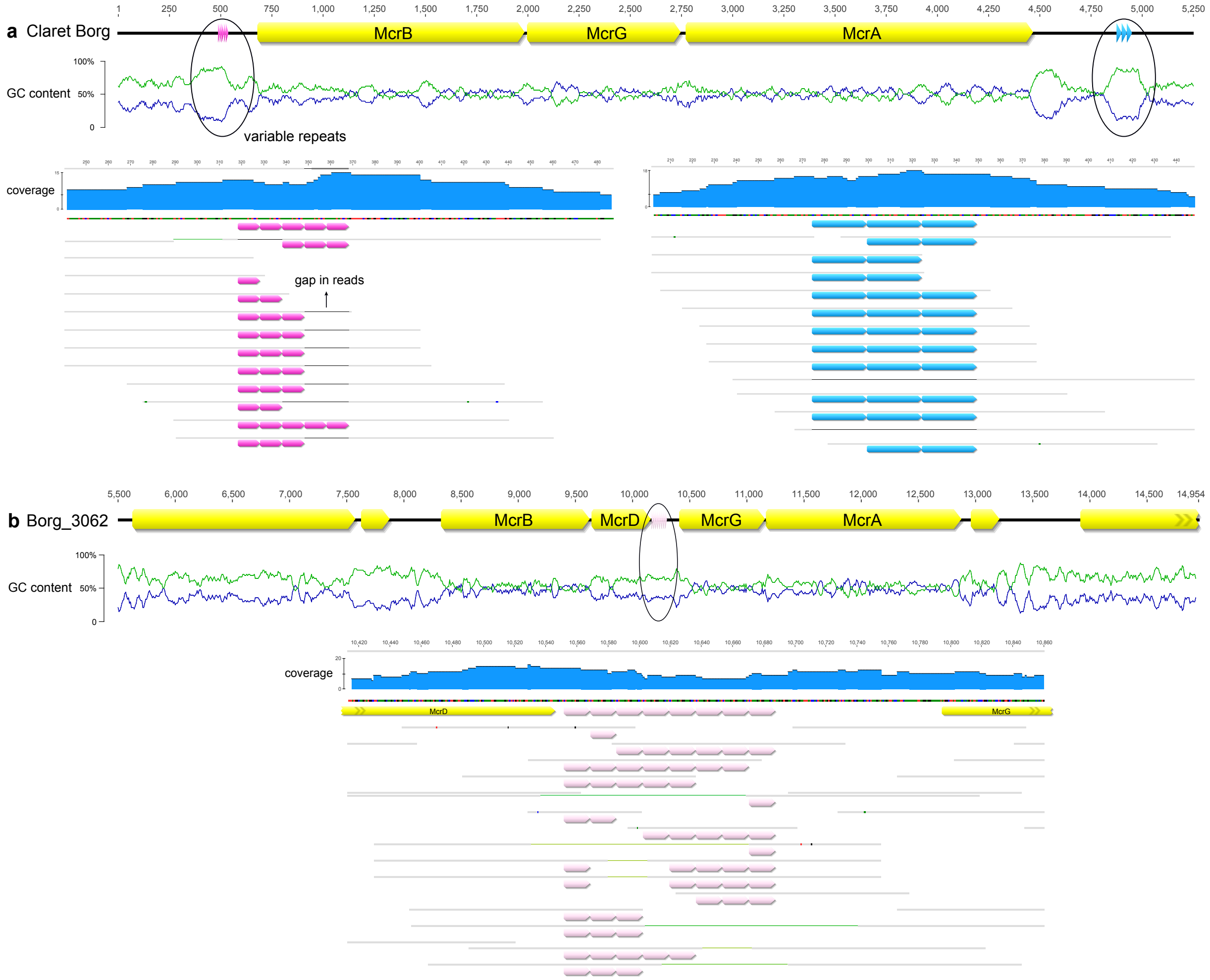


**Figure S9. Tandem repeats associated with *mcr* operons.** Read mapping shows the variable repeat numbers encoded flanking *mcr* operons (**a**) and between *mcrD* and *mcrG* (**b**) at low-GC regions (where green and blue lines indicate AT and GC contents, respectively). For example, some Borg strains encode five repeats at the upstream of *mcrB* (**a**), whereas some only encode two or three (pink arrows).

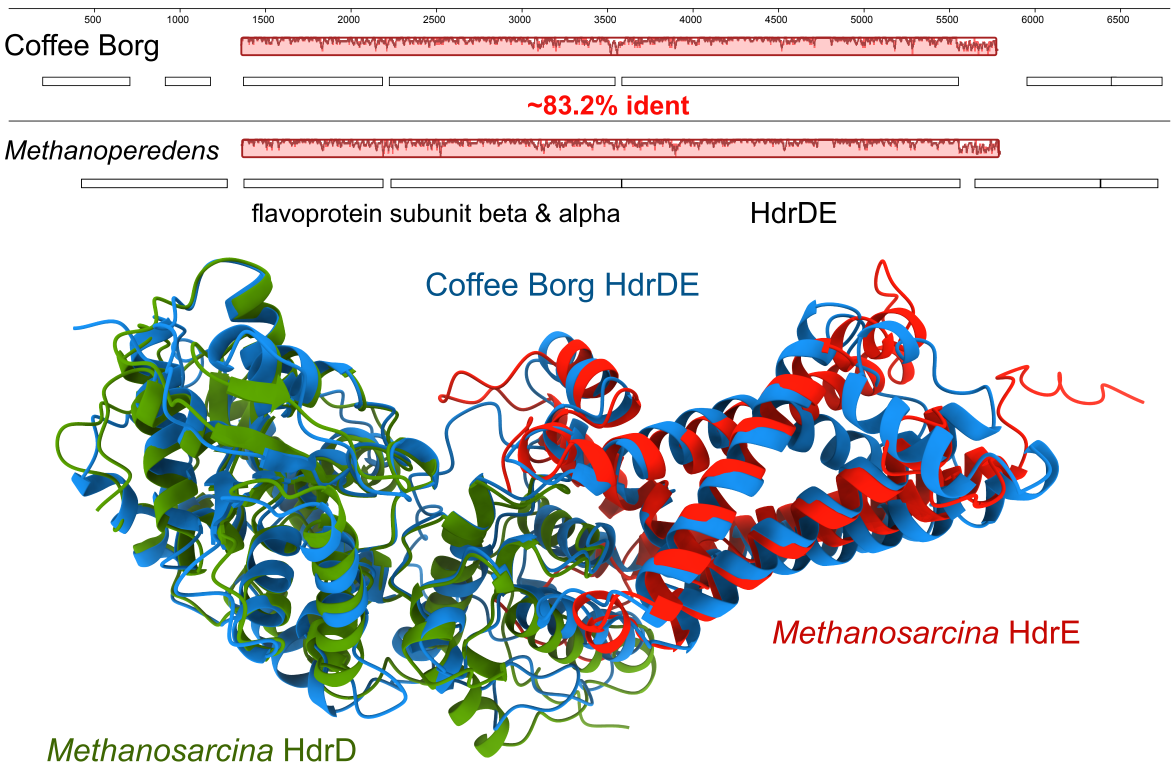


**Figure S10. Fused heterodisulfide reductase HdrDE encoded in Coffee Borg.** Genome regions of Coffee Borg and *Methanoperedens* were compared using Mauve alignment. The pairwise nucleotide identity of sequence blocks containing HdrDE and flavoproteins is ~83.2%. The structure of the Coffee HdrDE is predicted (blue) and superimposed with *Methanosarcina* HdrD (green; AF-P96797-F1-model_v6) and HdrE (red; AF-P96796-F1-model_v6). The TM scores are 0.88 and 0.69, respectively.

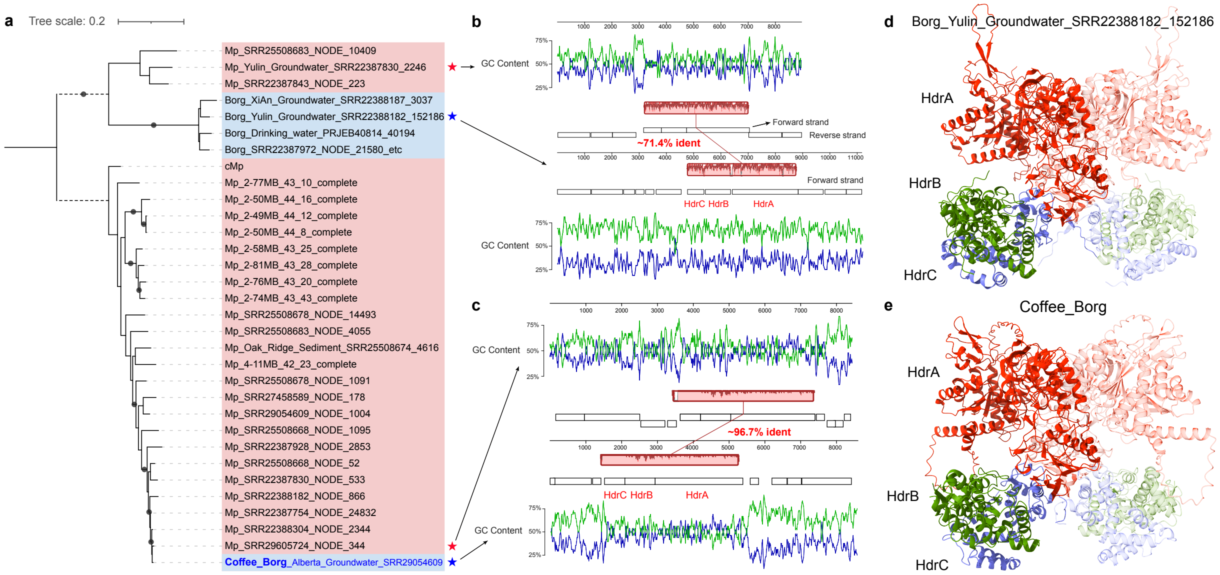


**Figure S11. Heterodisulfide reductase complex (HdrABC) encoded in Borgs. (a)** Phylogeny of HdrA proteins using the best-fit substitution model "Q.plant+R3". Sequences highlighted in red are from *Methanoperedens* and those in blue are from Borgs. Coffee Borg is a curated, complete genome. Branches are labeled when SH-aLRT values ≥ 80% and UFBoot values ≥ 95%. **(b-c)** Genomic comparison of *Methanoperedens* (labeled by red stars) and Borgs (labeled by blue stars) located in two phylogenetic clades. The *hdrABC* gene operons share high similarity, as depicted by Mauve alignment and pairwise nucleotide identity. Note that the *Methanoperedens* genomes have higher GC content than the Borgs (green and blue lines indicate AT and GC contents, respectively). Genes in the Borg are encoded unidirectionally, whereas in the *Methanoperedens* they are encoded in a mixed manner. **(d-e)** Predicted heterohexameric structures (A2B2C2) of HdrABC complexes encoded in Borgs. The ipTM and pTM values of the complex from a partial Borg genome are 0.81 and 0.8 **(c)**, and from Coffee Borg are 0.83 and 0.82 **(d)**. The best match of crystallized structures is the HdrABC complex from *Methanothermococcus thermolithotrophicus* (PDB: 5ODI; qTM=0.78). The delta subunit of methyl-viologen-reducing hydrogenase in *M. thermolithotrophicus* is fused to the C-terminal end of HdrA in Borgs and *Methanoperedens*.

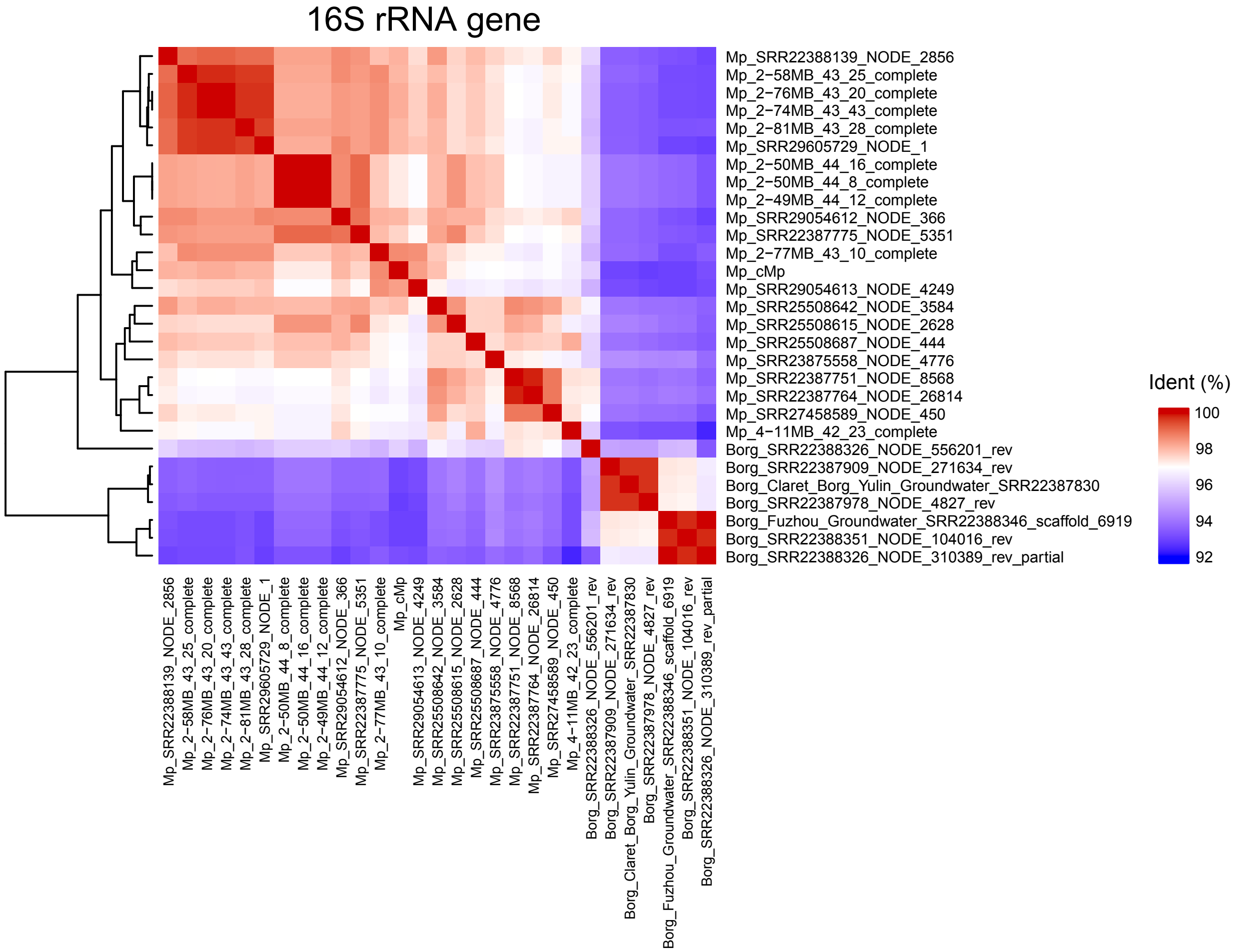


**Figure S12. Comparison of 16S rRNA genes encoded in Borgs and *Methanoperedens*.** Pairwise identity was calculated using blastn. Genomes are clustered based on Euclidean distances and complete linkage.

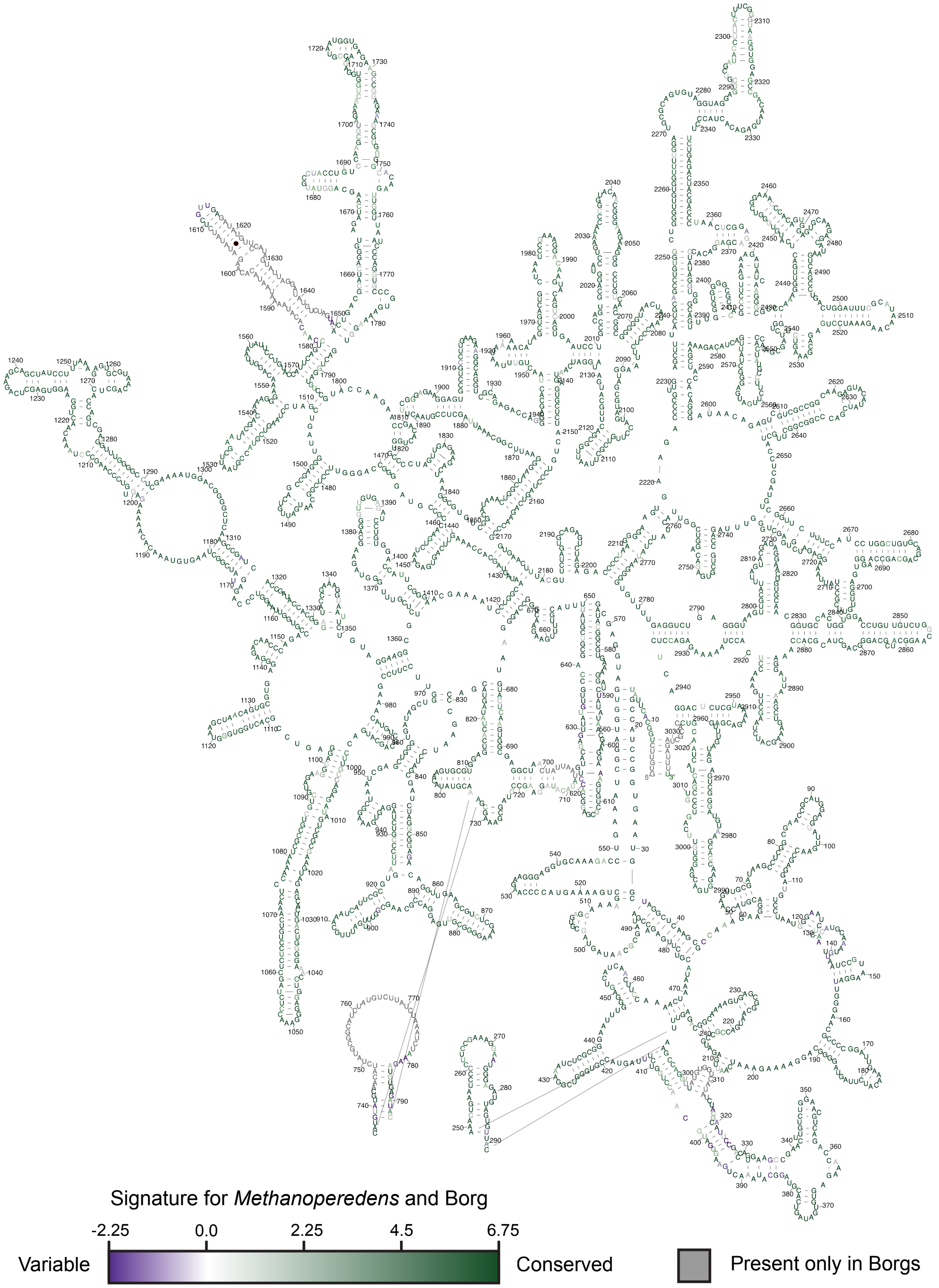


**Figure S13. 2D model of the Borg 23S rRNA.** Conservation between *Methanoperedens* and Borg sequences was calculated with TwinCons and is shown with a purple-green gradient with gray nucleotides marking Borg-only presence.

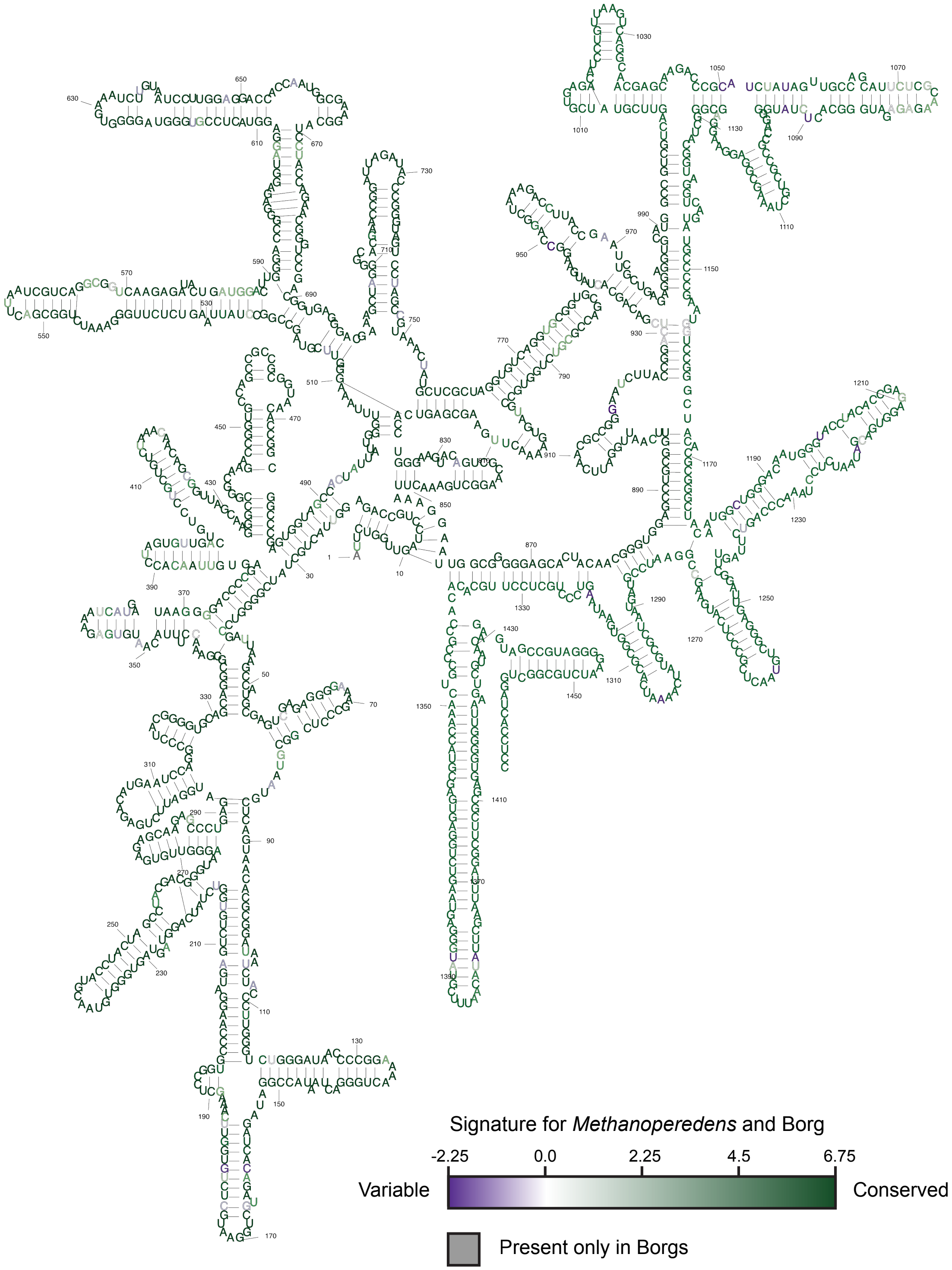


**Figure S14. 2D model of the Borg 16S rRNA.** Conservation between *Methanoperedens* and Borg sequences was calculated with TwinCons and is shown with a purple-green gradient with gray nucleotides marking Borg-only presence.

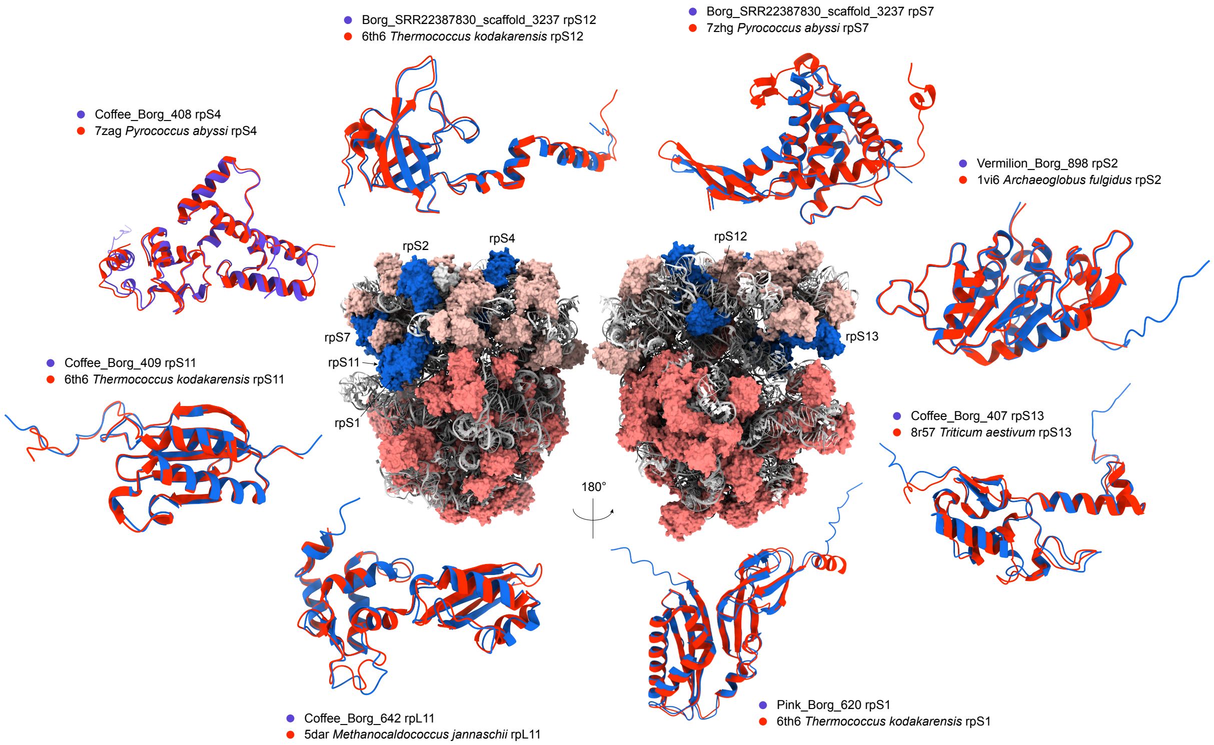


**Figure S15. Ribosomal proteins encoded in Borgs.** The 70S ribosome of *Thermococcus kodakarensis* (PDB: 6TH6) is shown in the middle with the ribosomal proteins in Borgs colored in blue, the remaining large ribosomal subunit proteins (rpL) in salmon, the small ribosomal subunit proteins (rpS) in light salmon, and the RNAs in gray cartoon. Each Borg ribosomal protein (blue) is structurally superimposed with the best match in the PDB database (red). Note that rpL11 is not included in the *Thermococcus kodakarensis* ribosome.

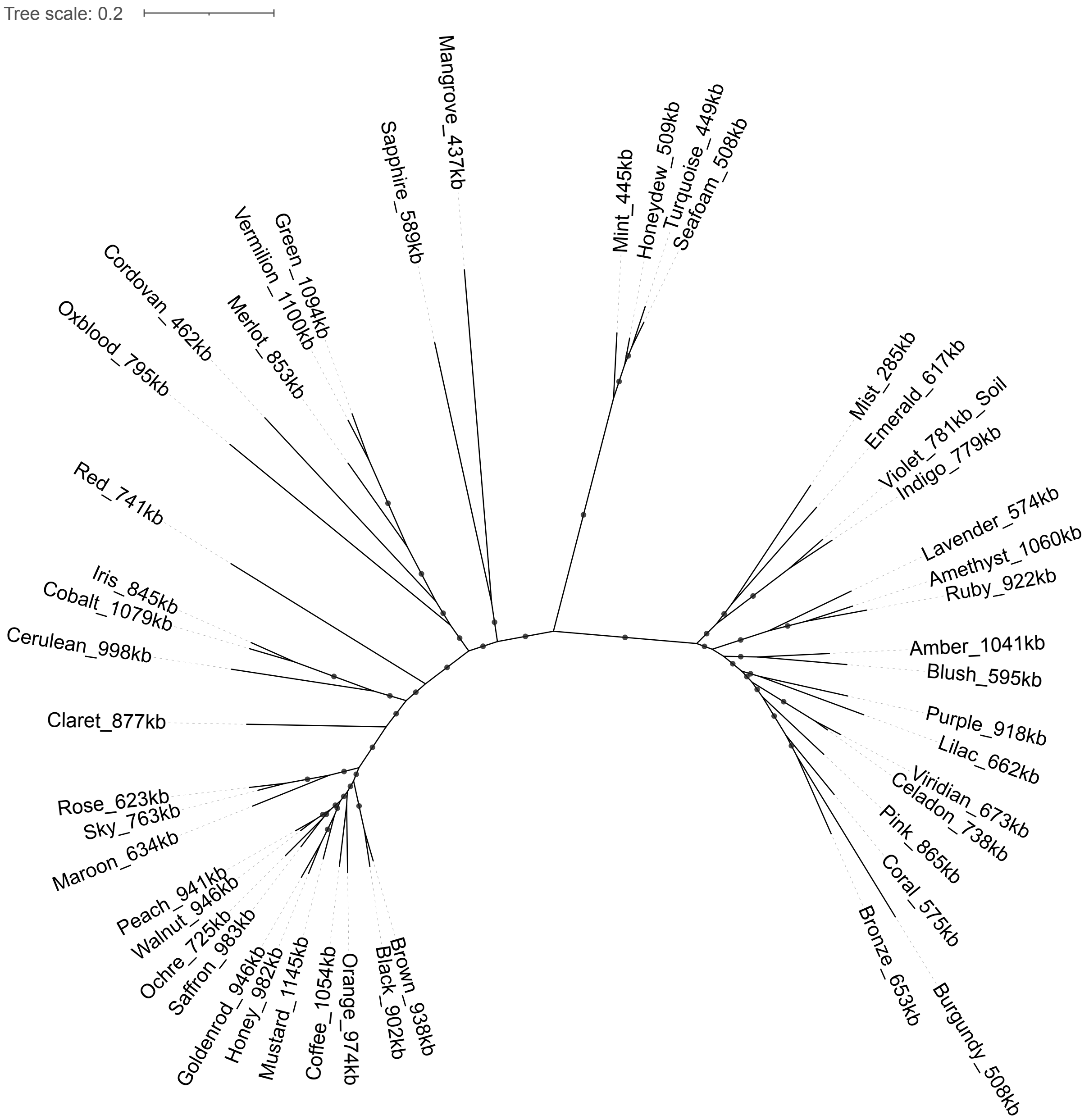


**Figure S16. Phylogeny of Borgs based on the concatenation of 45 single-copy proteins.** The alignment includes ~10,300 amino acids. The tree was constructed using the best-fit substitution model “Q.insect+F+R7”. Branches are labeled when SH-aLRT values ≥ 80% and UFBoot values ≥ 95%.

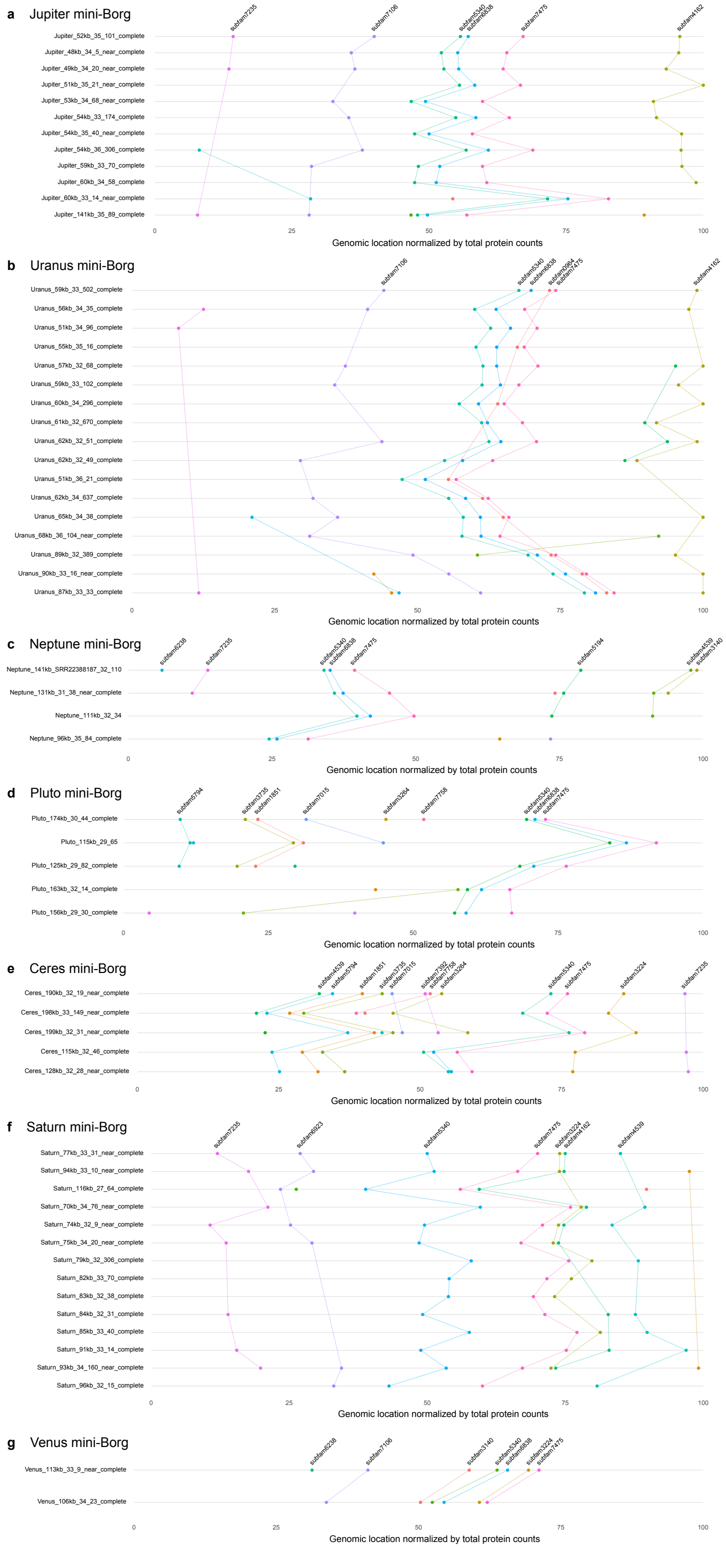


**Figure S17. Distribution and order of 45 single-copy genes within each mini-Borg clade.** Dots indicate the relative genomic locations of the genes normalized by total protein counts. Lines connect the same genes across genomes.

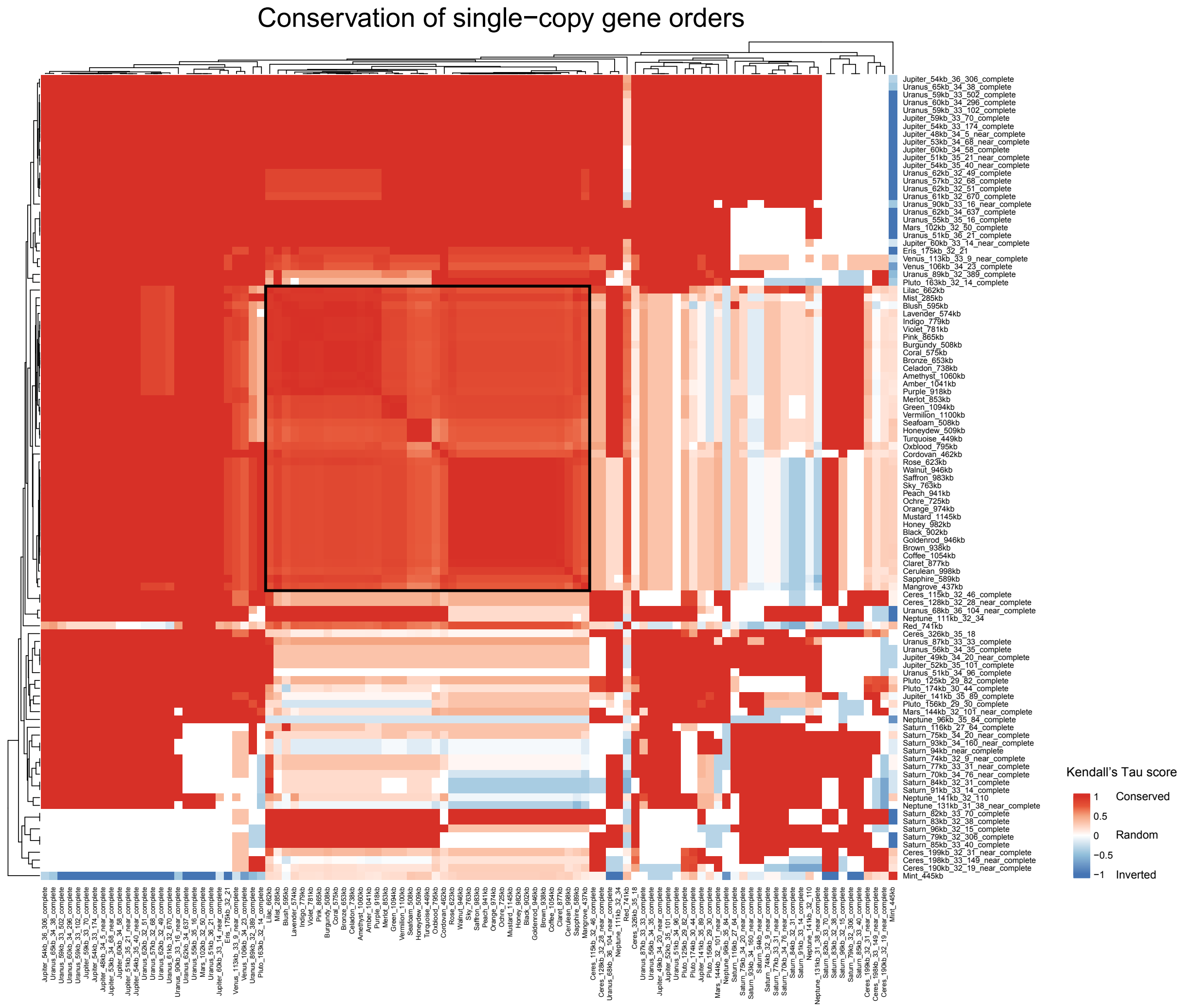


**Figure S18. Conservation of single-copy gene orders in Borgs and mini-Borgs.** Kendall Tau scores are calculated based on the relative genomic location of shared single-copy genes between genome pairs. Genomes are clustered by Euclidean distance (average linkage). According to the result within Borgs (outlined by a black square), a pairwise Kendall Tau score > 0.5 indicates that two genomes share significantly conserved orders of single-copy marker genes.

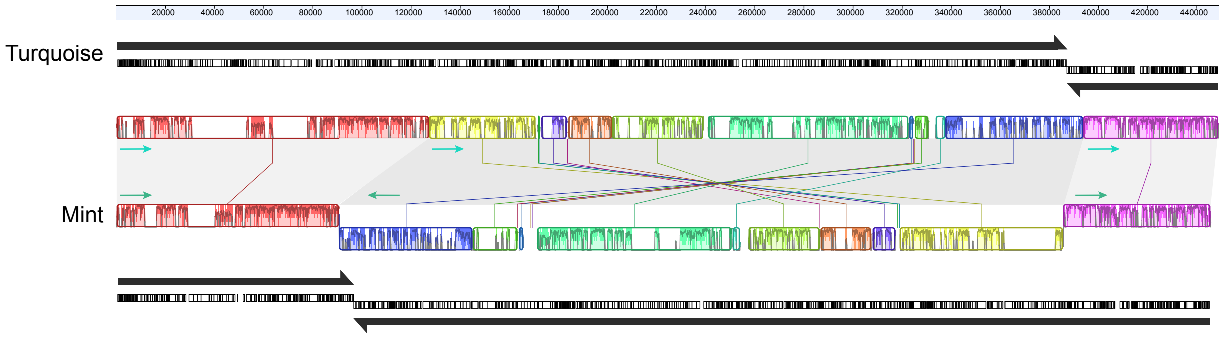


**Figure S19. Genome rearrangement in Borg genomes.** Turquoise Borg is aligned with the phylogenetically closed Mint using Mauve alignment. Long black arrows indicate genome replichores and coding strands. Note that the middle genome regions are aligned in opposite directions.

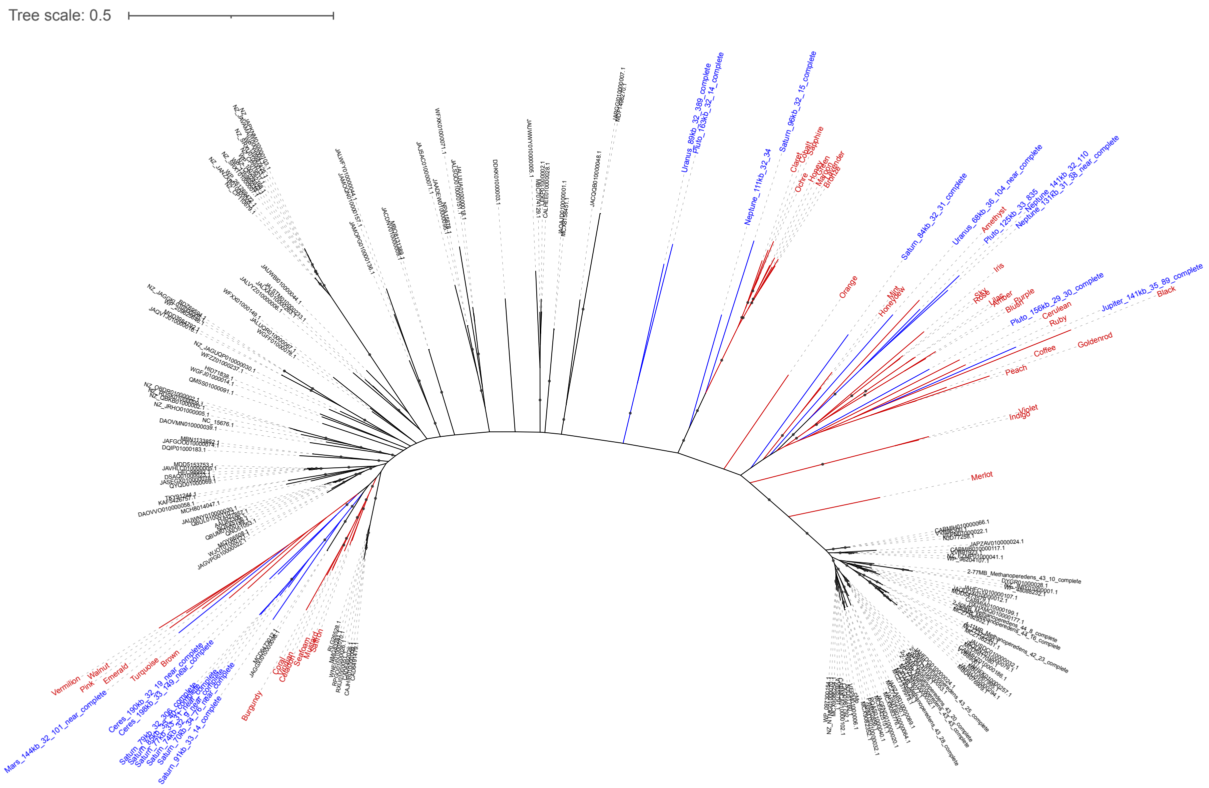


**Figure S20. Phylogeny of single-copy myo-inositol-1-phosphate synthases (subfam4539) encoded in Borgs and mini-Borgs.** Sequences (branches) in red and blue indicate Borg and mini-Borg genes, and those in black are non-Borg reference sequences. The tree was constructed using the best-fit substitution model “LG+F+I+R6”. Branches are labeled when SH-aLRT values ≥ 80% and UFBoot values ≥ 95%.

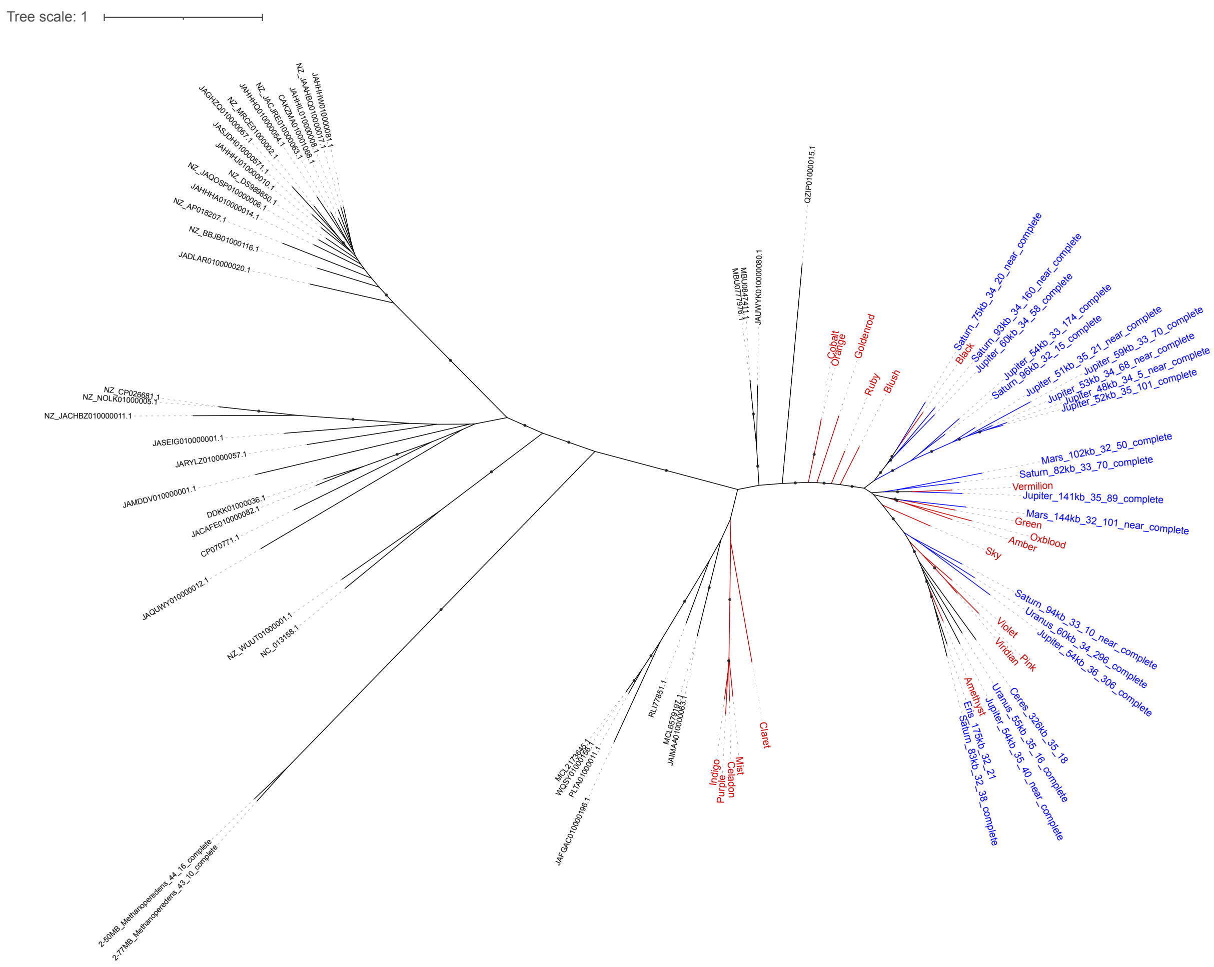


**Figure S21. Phylogeny of single-copy alpha-maltose-1-phosphate synthases (subfam4861) encoded in Borgs and mini-Borgs.** Sequences (branches) in red and blue indicate Borg and mini-Borg genes, and those in black are non-Borg reference sequences. The tree was constructed using the best-fit substitution model “LG+F+I+R6”. Branches are labeled when SH-aLRT values ≥ 80% and UFBoot values ≥ 95%.

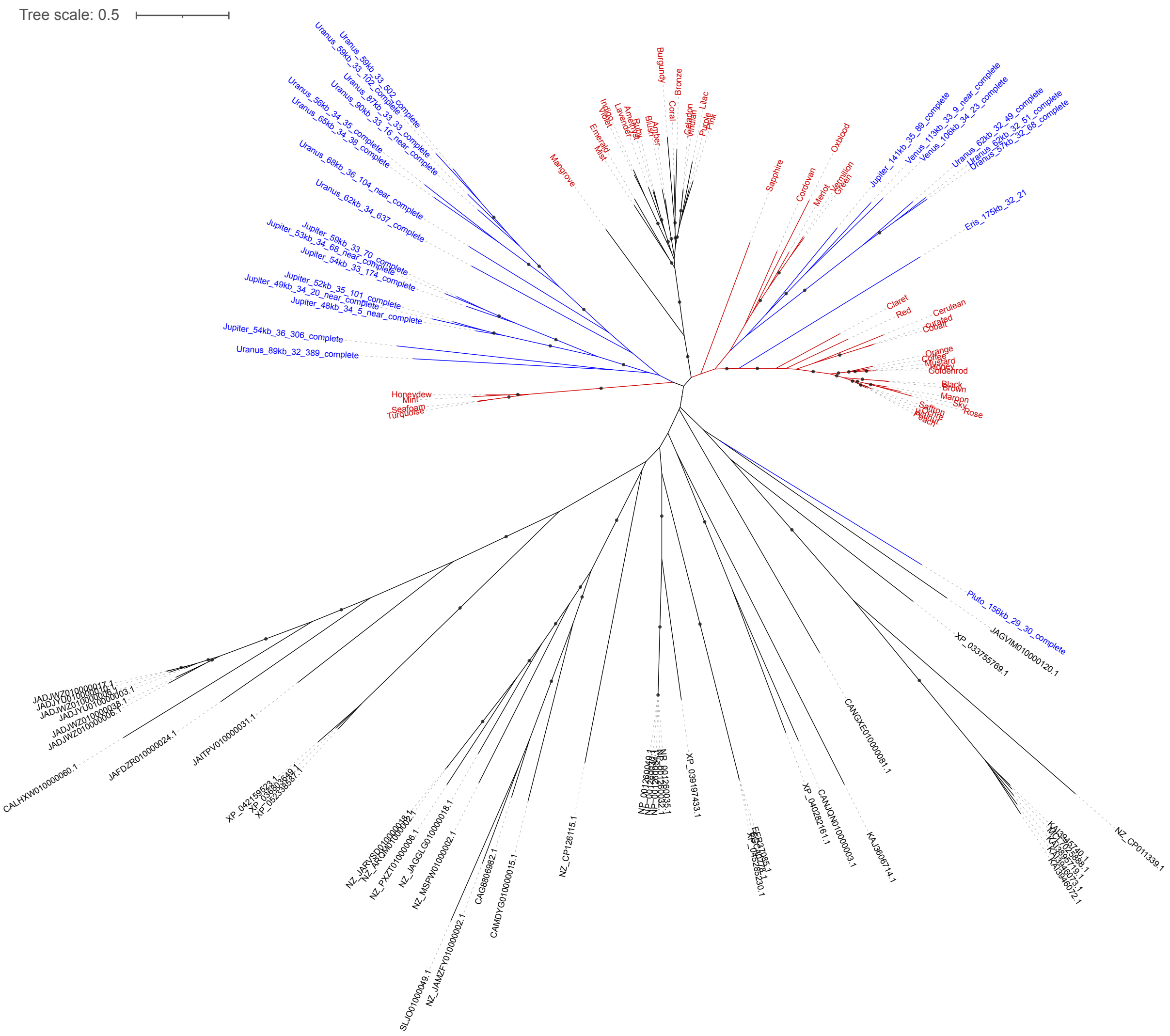


**Figure S22. Phylogeny of single-copy hypothetical proteins (subfam7106) encoded in Borgs and mini-Borgs.** Sequences (branches) in red and blue indicate Borg and mini-Borg genes, and those in black are non-Borg reference sequences. The tree was constructed using the best-fit substitution model “WAG+F+R5”. Branches are labeled when SH-aLRT values ≥ 80% and UFBoot values ≥ 95%.

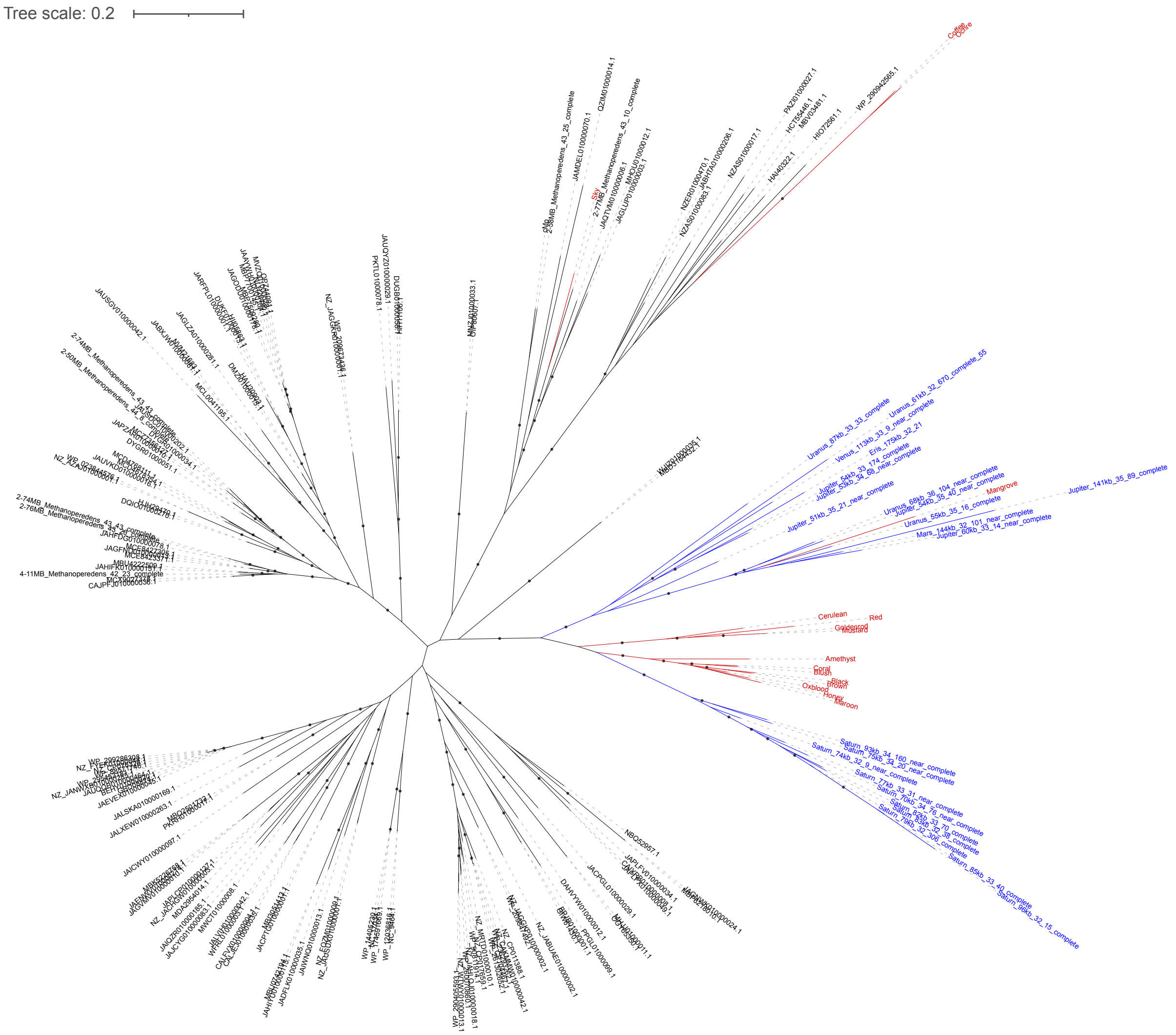


**Figure S23. Phylogeny of single-copy GDP-mannose 4,6-dehydratases (subfam4704) encoded in Borgs and mini-Borgs.** Sequences (branches) in red and blue indicate Borg and mini-Borg genes, and those in black are non-Borg reference sequences. The tree was constructed using the best-fit substitution model “LG+I+R5”. Branches are labeled when SH-aLRT values ≥ 80% and UFBoot values ≥ 95%.

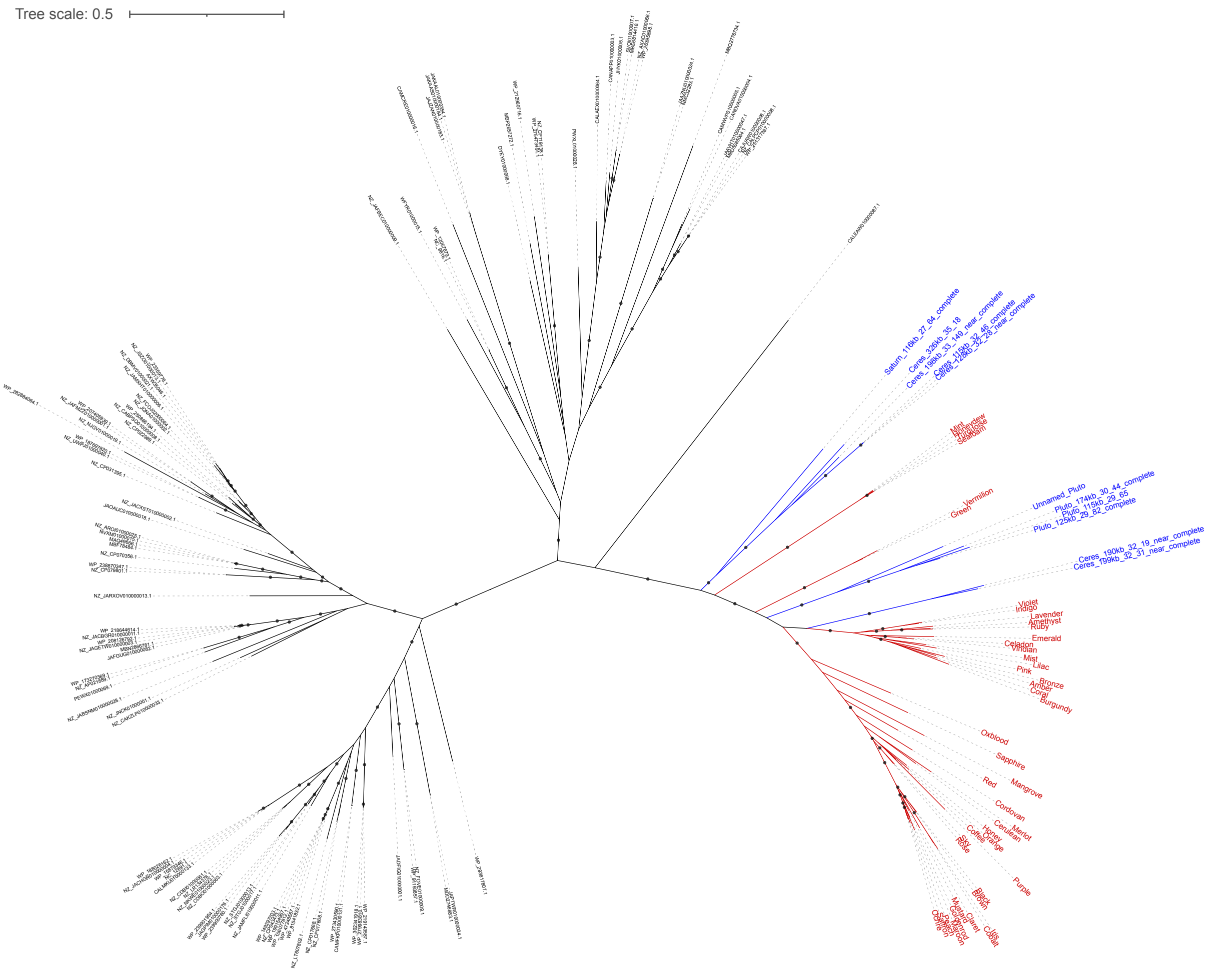


**Figure S24. Phylogeny of single-copy DNA recombinases (subfam1851) encoded in Borgs and mini-Borgs.** Sequences (branches) in red and blue indicate Borg and mini-Borg genes, and those in black are non-Borg reference sequences. The tree was constructed using the best-fit substitution model “LG+I+R5”. Branches are labeled when SH-aLRT values ≥ 80% and UFBoot values ≥ 95%.

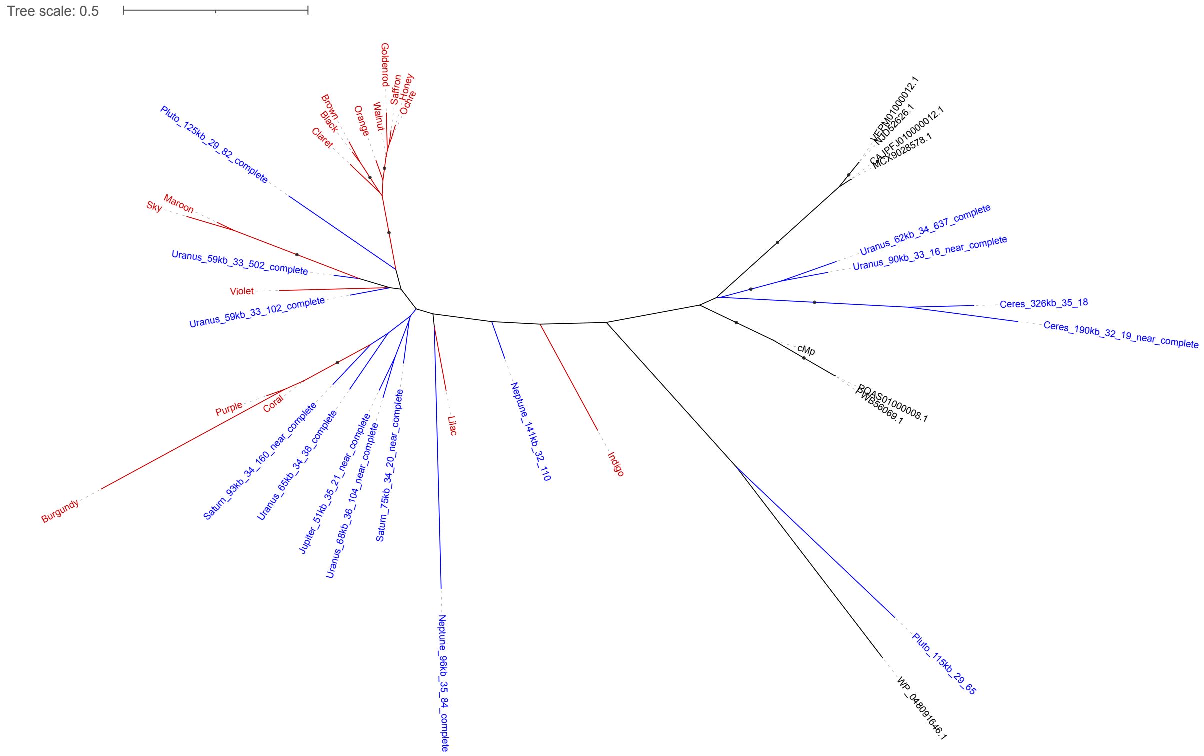


**Figure S25. Phylogeny of single-copy hypothetical proteins (subfam2870) encoded in Borgs and mini-Borgs.** Sequences (branches) in red and blue indicate Borg and mini-Borg genes, and those in black are non-Borg reference sequences. The tree was constructed using the best-fit substitution model “Q.insect+G4”. Branches are labeled when SH-aLRT values ≥ 80% and UFBoot values ≥ 95%.

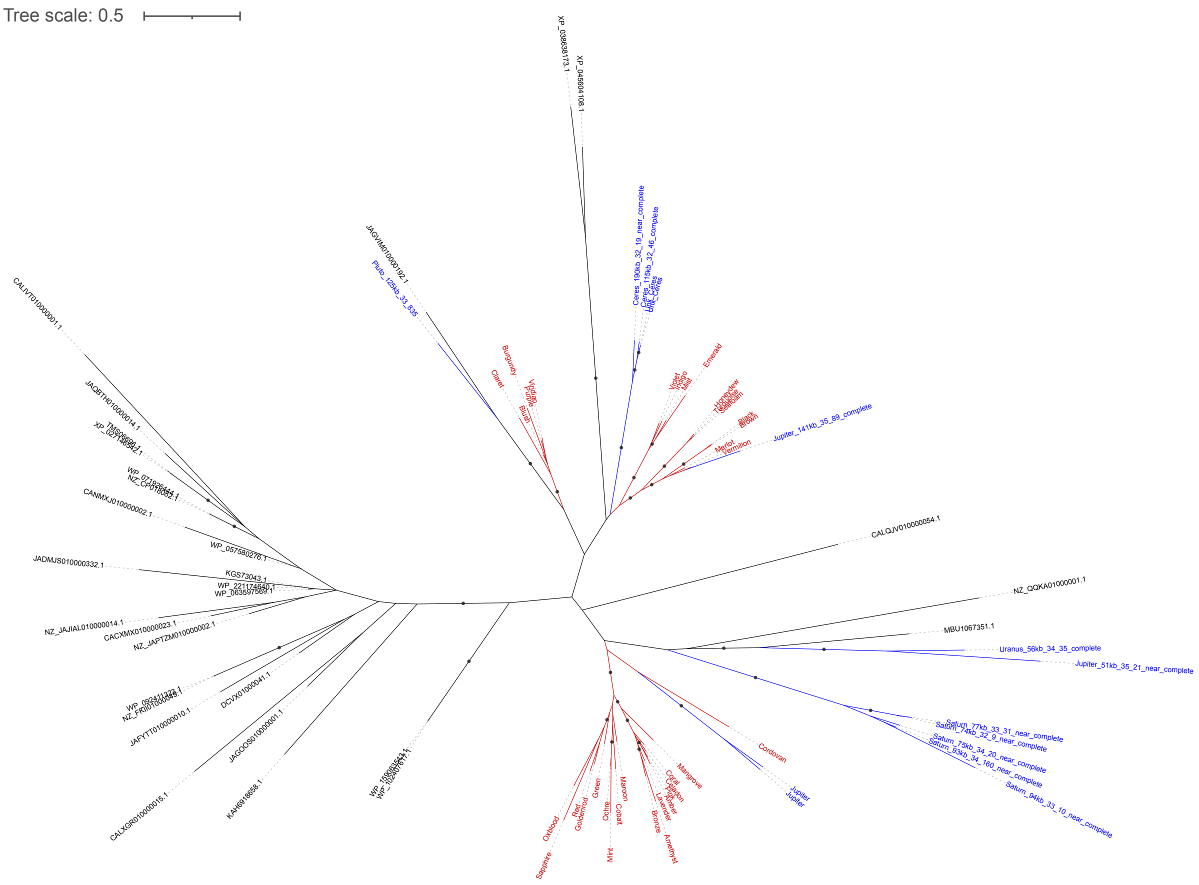


**Figure S26. Phylogeny of single-copy hypothetical proteins (subfam4208) encoded in Borgs and mini-Borgs.** Sequences (branches) in red and blue indicate Borg and mini-Borg genes, and those in black are non-Borg reference sequences. The tree was constructed using the best-fit substitution model “VT+F+G4”. Branches are labeled when SH-aLRT values ≥ 80% and UFBoot values ≥ 95%.

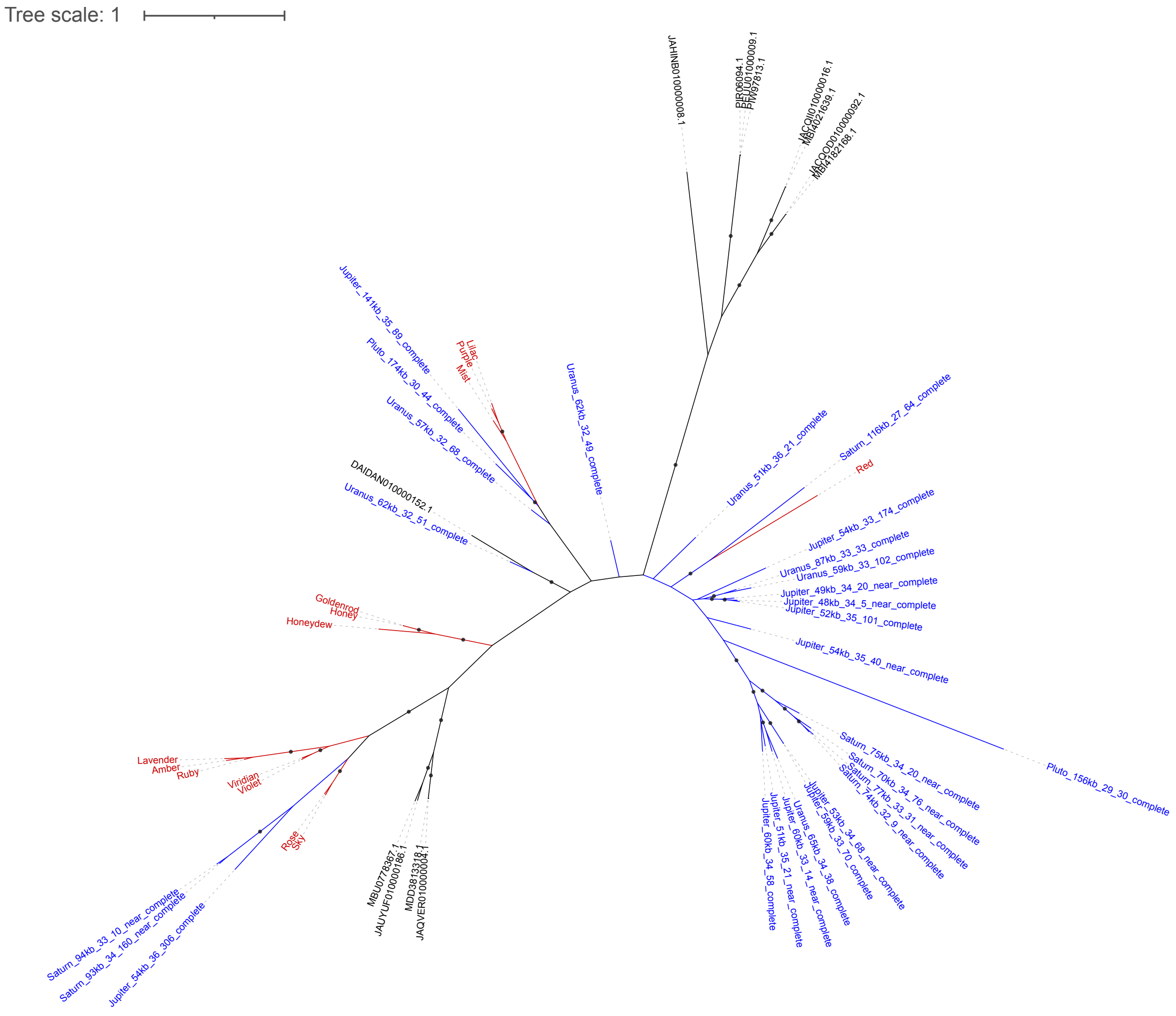


**Figure S27. Phylogeny of single-copy hypothetical proteins (subfam5145) encoded in Borgs and mini-Borgs.** Sequences (branches) in red and blue indicate Borg and mini-Borg genes, and those in black are non-Borg reference sequences. The tree was constructed using the best-fit substitution model “WAG+F+R4”. Branches are labeled when SH-aLRT values ≥ 80% and UFBoot values ≥ 95%.

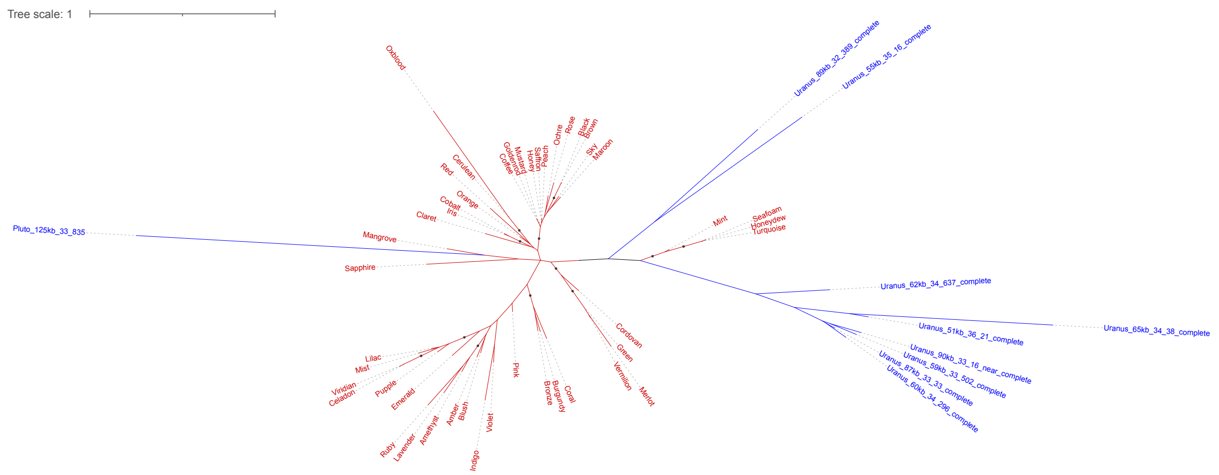


**Figure S28. Phylogeny of single-copy hypothetical proteins (subfam0964) encoded in Borgs and mini-Borgs.** Sequences (branches) in red and blue indicate Borg and mini-Borg genes. No homologs are found in other elements. The tree was constructed using the best-fit substitution model "Q.yeast+I+G4". Branches are labeled when SH-aLRT values ≥ 80% and UFBoot values ≥ 95%.

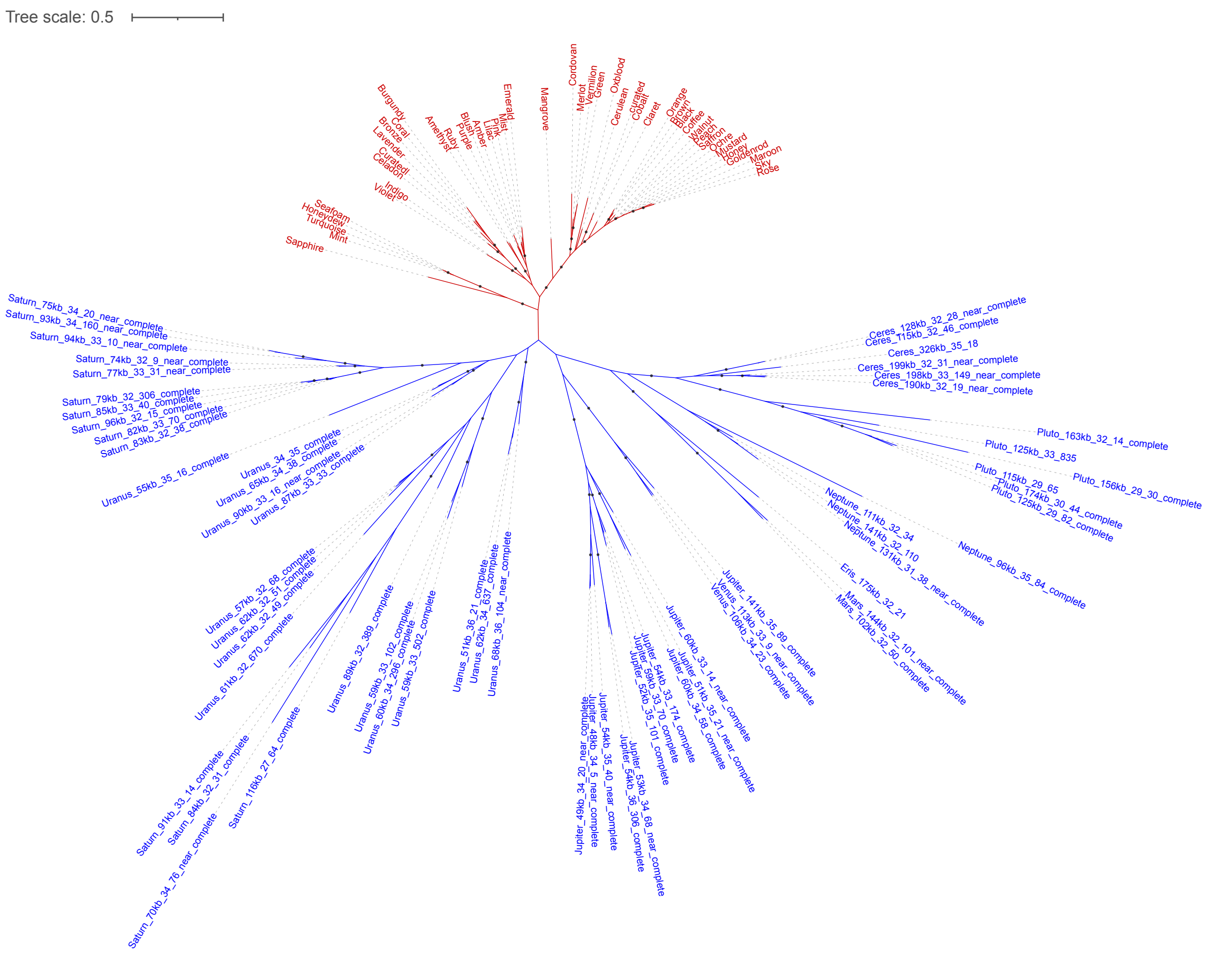


**Figure S29. Phylogeny of single-copy hypothetical proteins (subfam5340) encoded in Borgs and mini-Borgs.** Sequences (branches) in red and blue indicate Borg and mini-Borg genes. No homologs are found in other elements. The tree was constructed using the best-fit substitution model “Q.yeast+I+R5”. Branches are labeled when SH-aLRT values ≥ 80% and UFBoot values ≥ 95%.

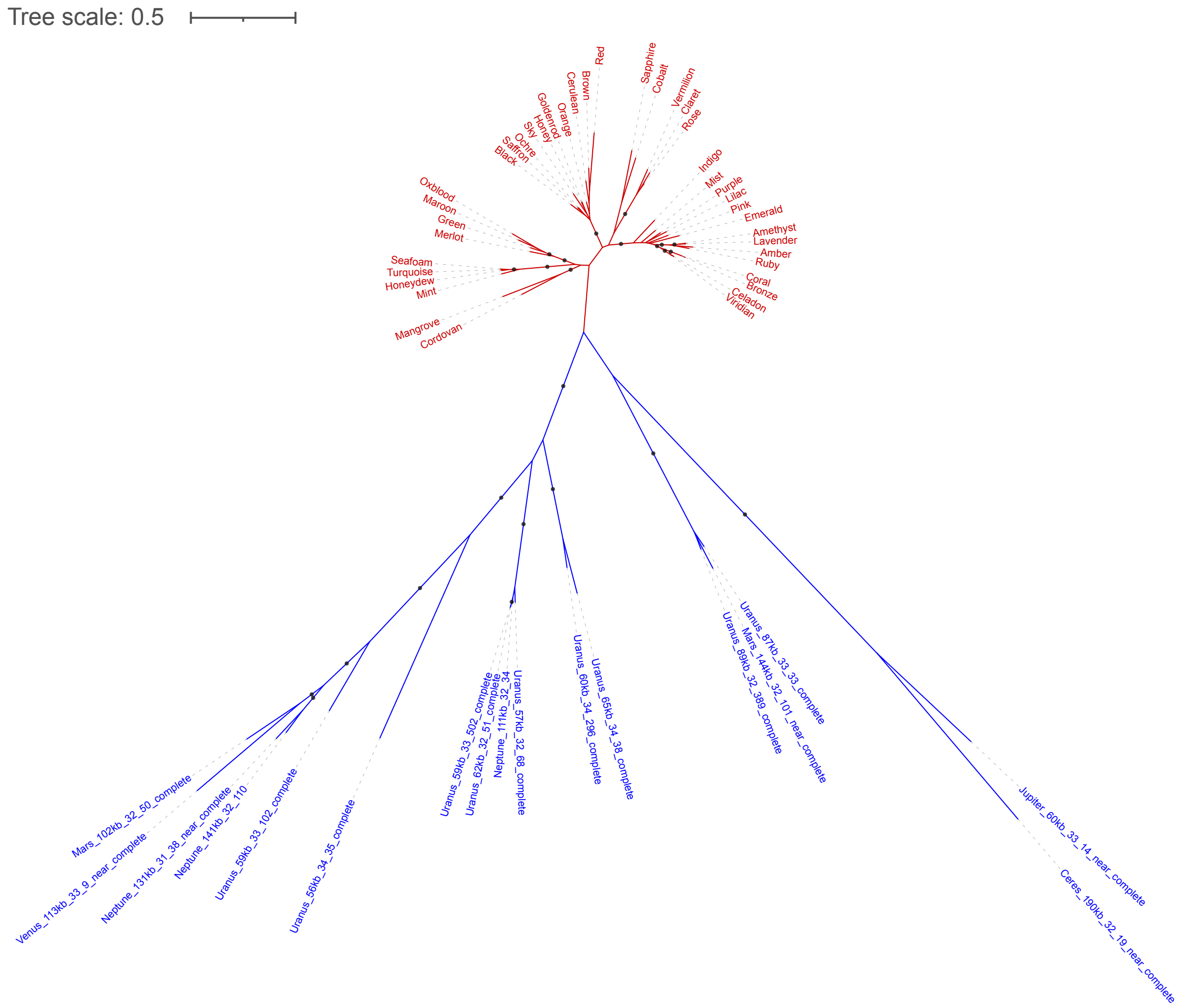


**Figure S30. Phylogeny of single-copy ATP-binding proteins (subfam1112) encoded in Borgs and mini-Borgs.** Sequences (branches) in red and blue indicate Borg and mini-Borg genes. The tree was constructed using the best-fit substitution model "Q.yeast+F+R4". Branches are labeled when SH-aLRT values ≥ 80% and UFBoot values ≥ 95%.

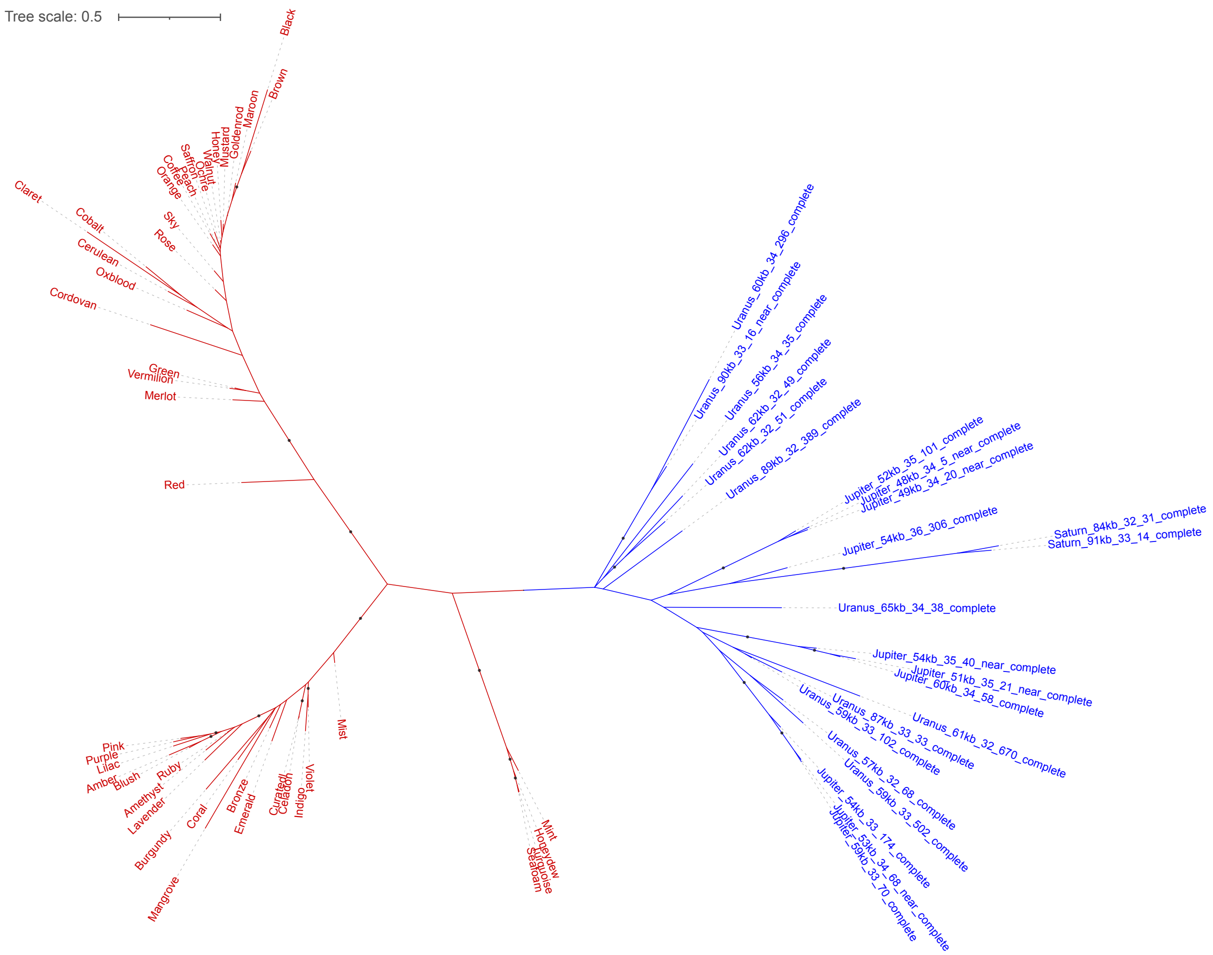


**Figure S31. Phylogeny of single-copy hypothetical proteins (subfam4162) encoded in Borgs and mini-Borgs.** Sequences (branches) in red and blue indicate Borg and mini-Borg genes. No homologs are found in other elements. The tree was constructed using the best-fit substitution model “Q.yeast+I+G4”. Branches are labeled when SH-aLRT values ≥ 80% and UFBoot values ≥ 95%.

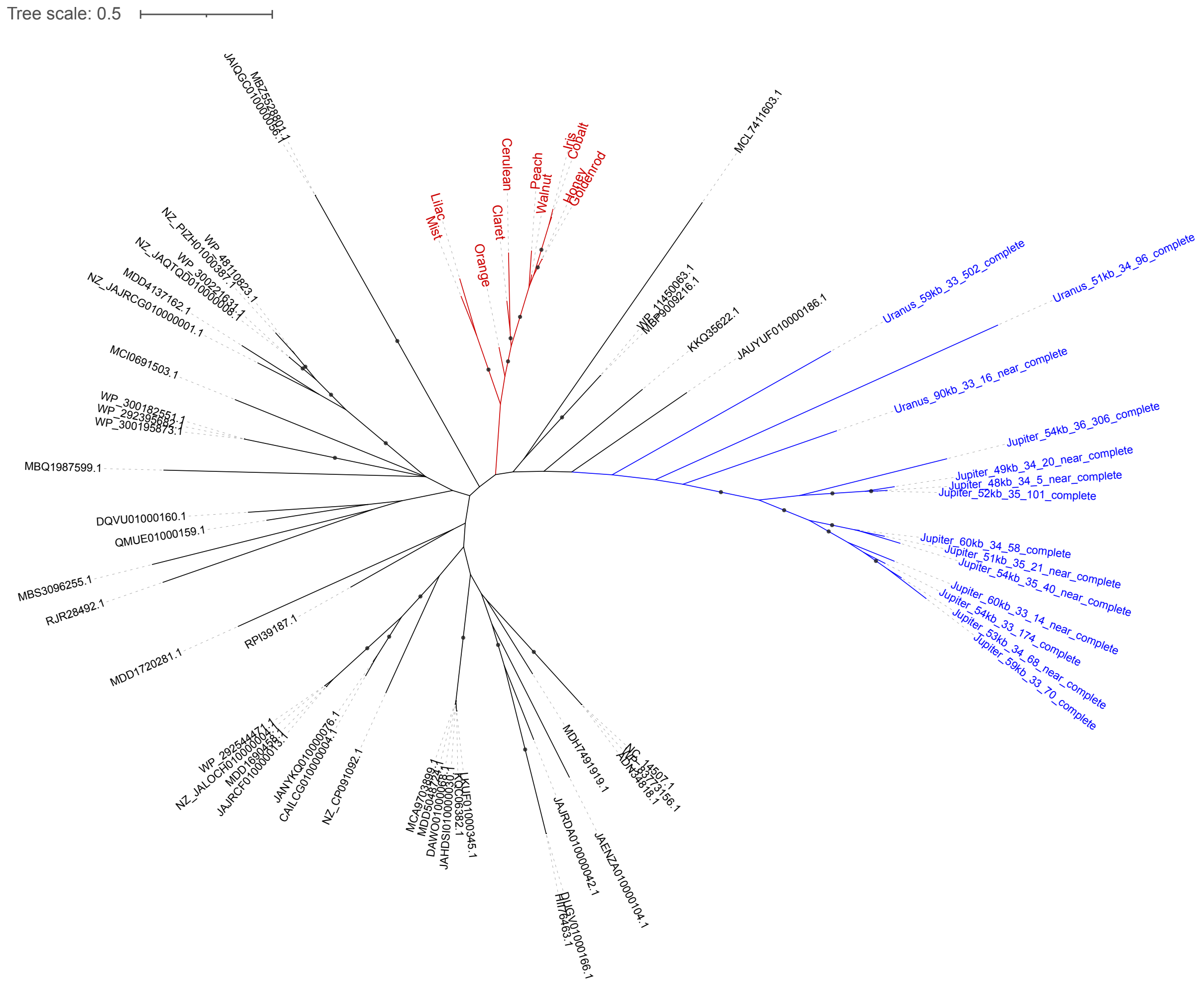


**Figure S32. Phylogeny of single-copy PEGA domain proteins (subfam5350) encoded in Borgs and mini-Borgs.** Sequences (branches) in red and blue indicate Borg and mini-Borg genes, and those in black are non-Borg reference sequences. The tree was constructed using the best-fit substitution model “Q.yeast+R3”. Branches are labeled when SH-aLRT values ≥ 80% and UFBoot values ≥ 95%.

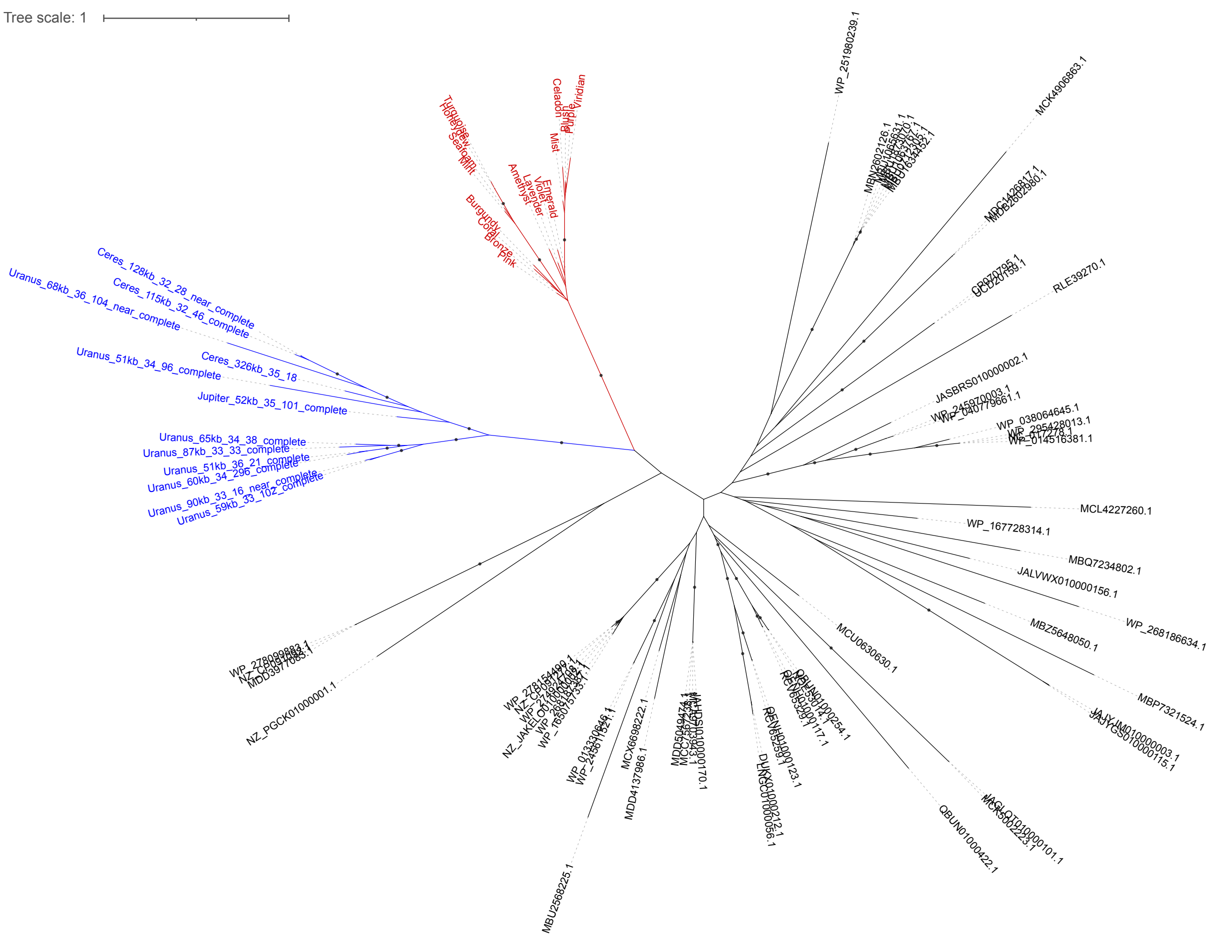


**Figure S33. Phylogeny of single-copy PEGA domain proteins (subfam6889) encoded in Borgs and mini-Borgs.** Sequences (branches) in red and blue indicate Borg and mini-Borg genes, and those in black are non-Borg reference sequences. The tree was constructed using the best-fit substitution model “Q.pfam+R4”. Branches are labeled when SH-aLRT values ≥ 80% and UFBoot values ≥ 95%.

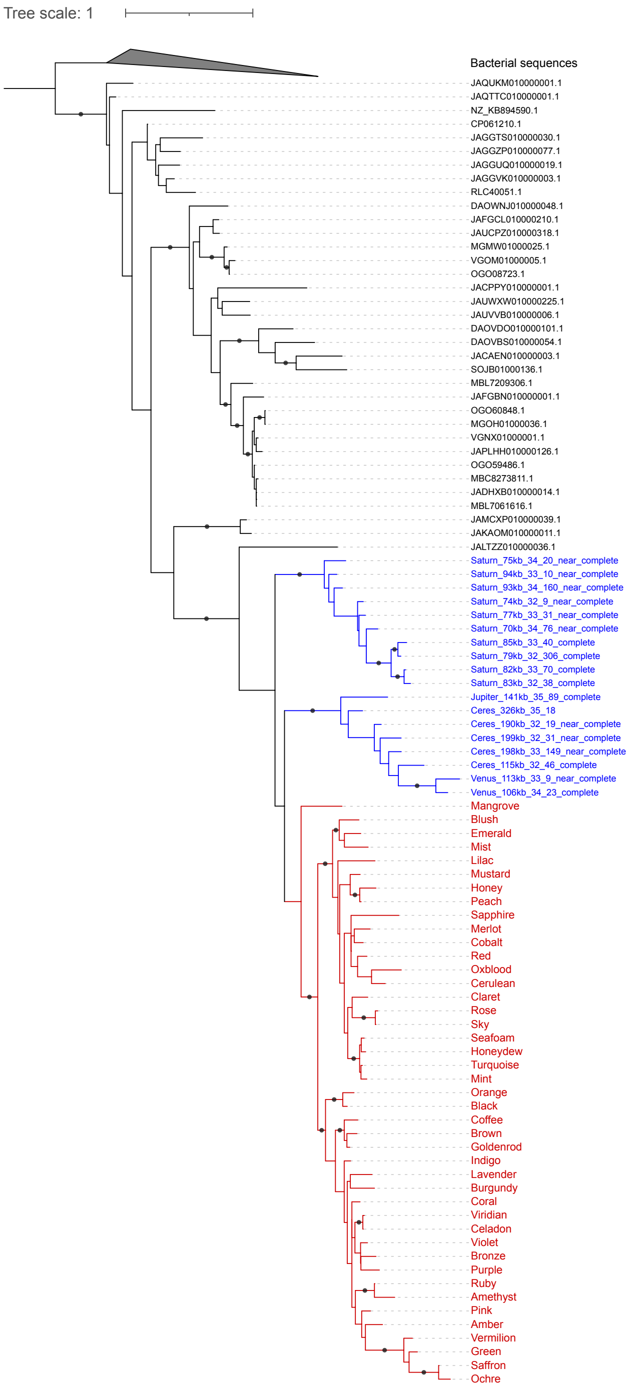


**Figure S34. Phylogeny of single-copy ferritins (subfam3224) encoded in Borgs and mini-Borgs.** Sequences (branches) in red and blue indicate Borg and mini-Borg genes, and those in black are non-Borg reference sequences. The tree was constructed using the best-fit substitution model “VT+R5” and rooted using bacterial sequences as the outgroup. Branches are labeled when SH-aLRT values ≥ 80% and UFBoot values ≥ 95%. Note that 44 of 47 Borgs encode this protein.

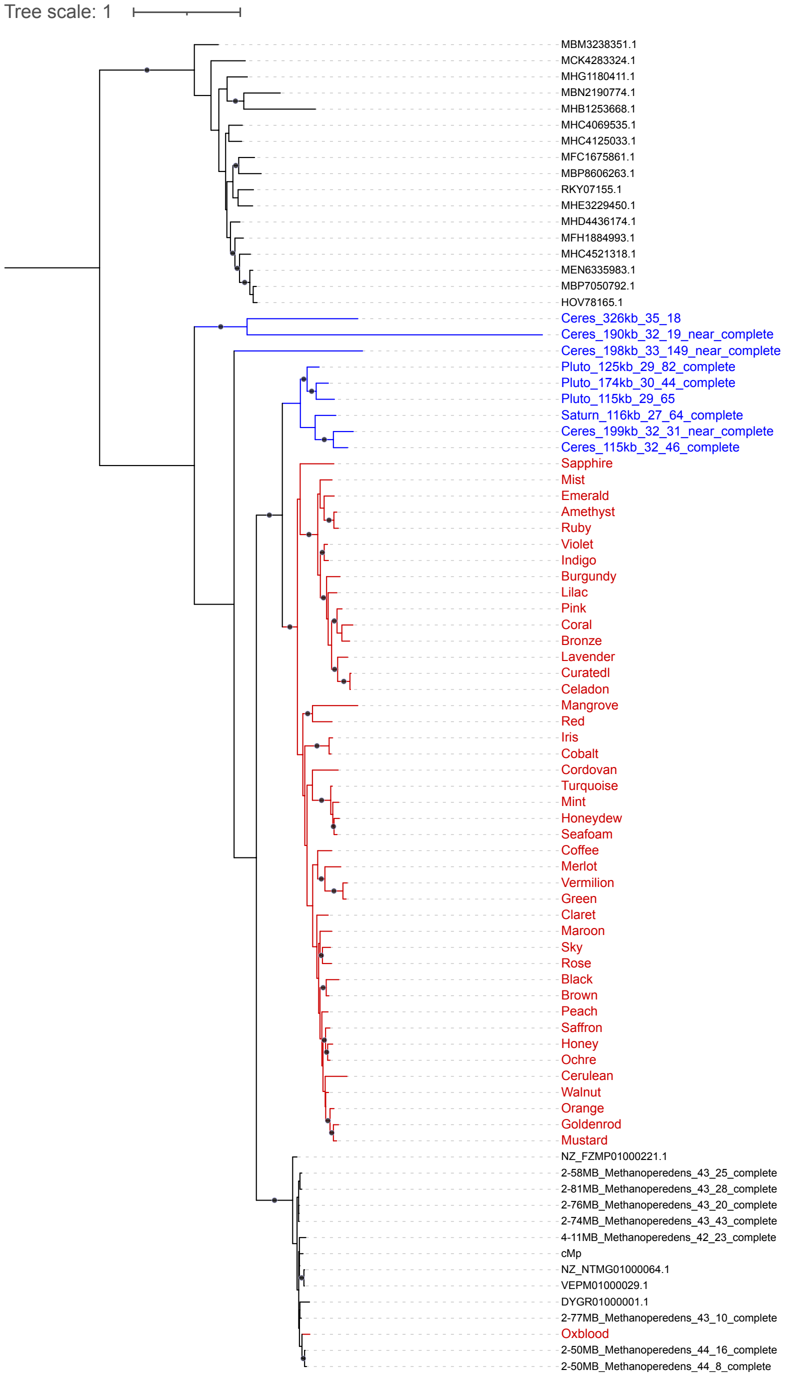


**Figure S35. Phylogeny of single-copy thymidylate synthases (subfam3735) encoded in Borgs and mini-Borgs.** Sequences (branches) in red and blue indicate Borg and mini-Borg genes, and those in black are non-Borg reference sequences. The tree was constructed using the best-fit substitution model "Q.insect+I+G4" and rooted using bacterial sequences as the outgroup. Branches are labeled when SH-aLRT values ≥ 80% and UFBoot values ≥ 95%. Note that 44 of 47 Borgs encode this protein.

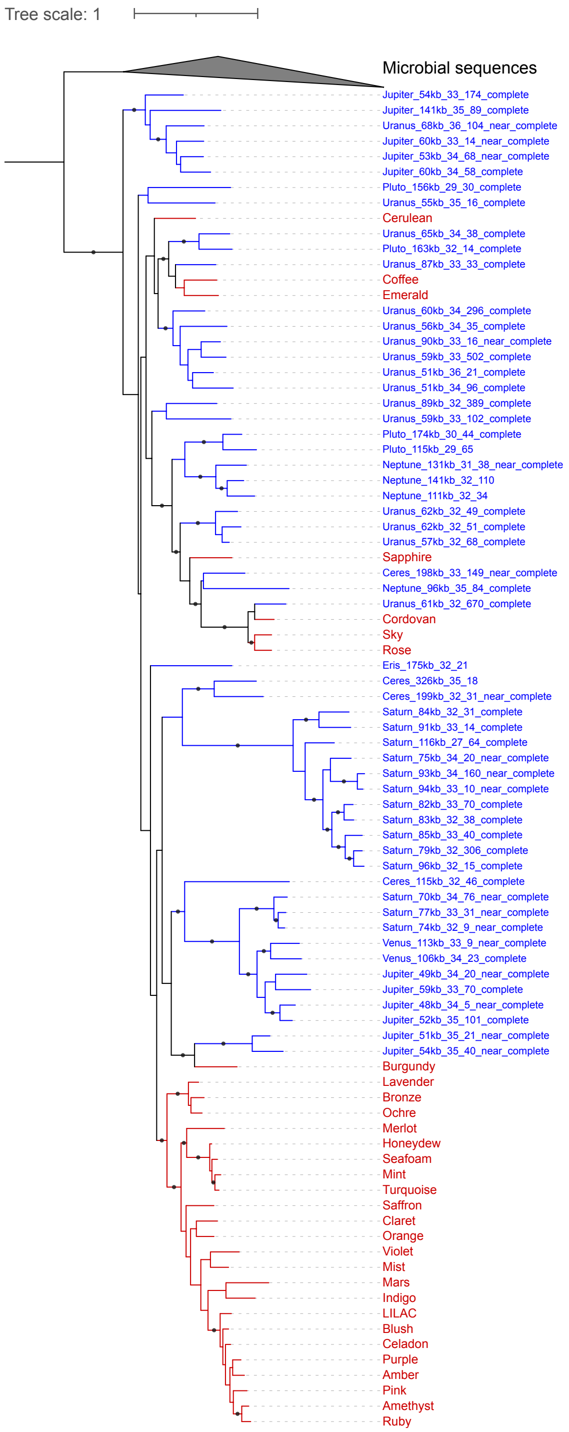


**Figure S36. Phylogeny of single-copy glycosyltransferases (subfam6077) encoded in Borgs and mini-Borgs.** Sequences (branches) in red and blue indicate Borg and mini-Borg genes. The tree was constructed using the best-fit substitution model “Q.pfam+F+R7” and rooted using microbial sequences as the outgroup. Branches are labeled when SH-aLRT values ≥ 80% and UFBoot values ≥ 95%.

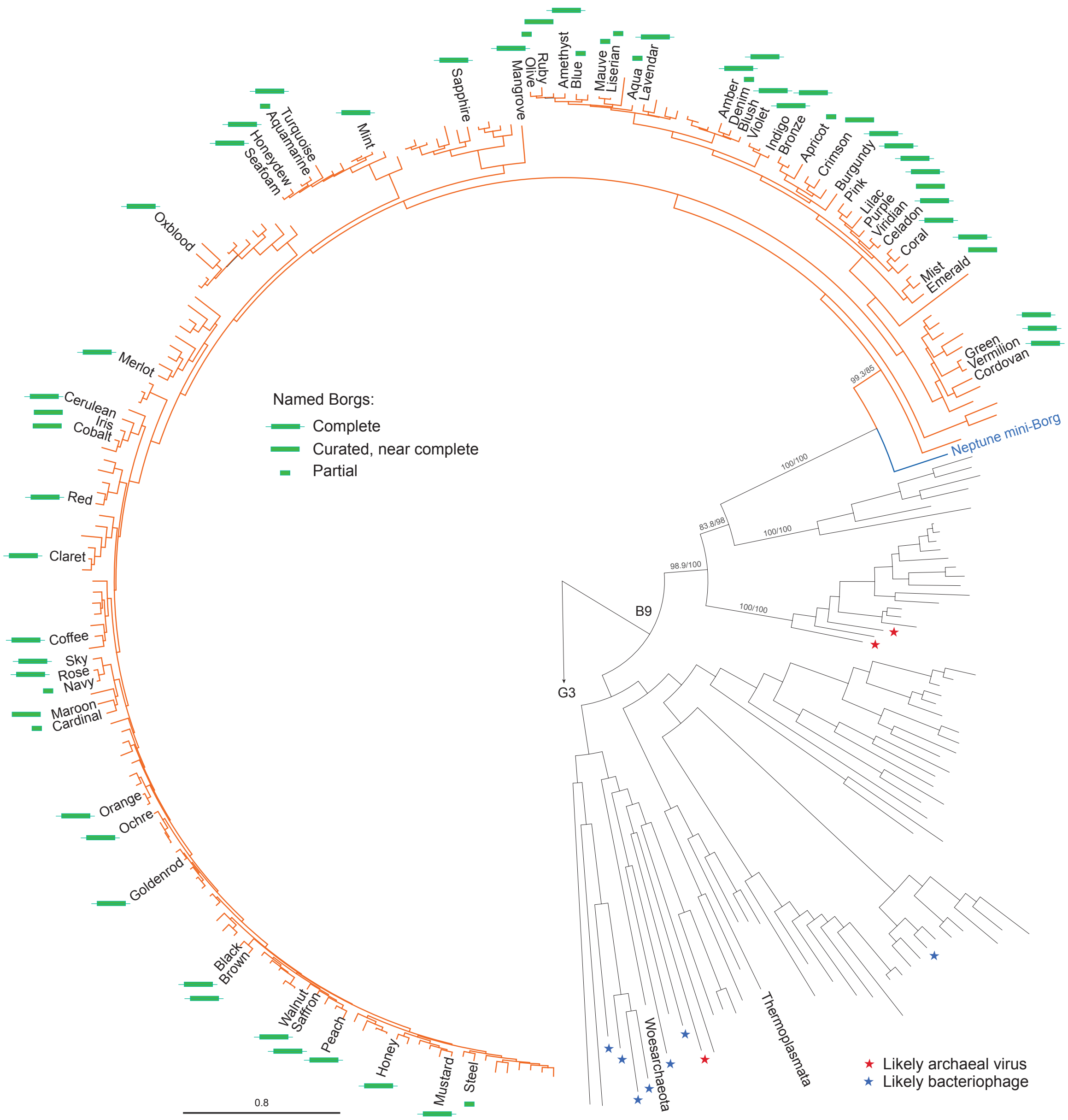


**Figure S37. DNA polymerase B9 encoded in Borgs and a Neptune mini-Borg.** Sequences (branches) in orange and purple indicate Borg and mini-Borg genes, and those in black are non-Borg reference sequences. The ML tree was constructed using the best-fit substitution model "LG+F+R8" and rooted using B-family DNA polymerase group G3 sequences as the outgroup. SH-aLRT and UFBoot values were calculated based on 1000 replicates and labeled on the major B9 clade branches.
